# Molecular Star Gazing: Development and Validation of an Environmental DNA Assay for the Imperiled Sunflower Sea Star (*Pycnopodia helianthoides*)

**DOI:** 10.64898/2026.05.07.723600

**Authors:** Zachary Gold, Kristin M. Robinson, Alyssa-Lois M. Gehman, Meghan M. Shea, Matthew A. Lemay, Johannes Weinrich, Colleen T.E. Kellogg, Rute B.G. Clemente-Carvalho, Lauren M. Schiebelhut, Alexandria B. Boehm, Ashley Kidd, Andrew Kim, Jason Hodin, Michael Dawson, Sean M. McAllister

## Abstract

The sunflower sea star (*Pycnopodia helianthoides*) suffered a catastrophic population decline across its range from 2013 to 2017 due to the devastating *Vibrio pectenicida* FHCF-3 driven sea star wasting disease (SSWD) pandemic with minimal signs of population recovery. The functional extinction of this apex predator across substantial parts of its range has created a need to identify and track the remaining intact populations. Environmental DNA (eDNA) approaches provide a simple, cost-effective, and non-destructive method for monitoring occurrences, and in some cases abundances, of marine species, consistently outperforming visual occurrence monitoring efforts in sensitivity, speed, and cost. Here, we designed, developed, and validated a *P. helianthoides*-specific eDNA assay to identify refugia, using both quantitative and digital droplet PCR approaches. We first generated the most comprehensive sea star mitochondrial genome reference database to date (n=93 taxa, n= 15 novel). We then used *unikseq* and Geneious bioinformatics software to identify the unique *nad5* gene region and design a highly specific hydrolysis probe-based PCR assay. We validated the performance of this assay through laboratory, mesocosm, and field testing, demonstrating a highly specific and sensitive assay. In a field application of the new assay across regions in British Columbia, Canada, we found a positive correlation between *P. helianthoides* eDNA concentrations and biomass density, especially when appropriately accounting for spatiotemporal integration scales (R^2^=0.67). The eDNA assay provides a rapid and scalable tool for monitoring the sunflower sea star which has been proposed for listing as threatened under the U.S. Endangered Species Act of 1973. Molecular tools like the one presented here enhance management and recovery efforts not only by identification and monitoring of remnant wild populations, but also by helping to assess population level response and recovery following reintroduction efforts.

## Introduction

Sunflower sea stars (*Pycnopodia helianthoides*) are apex predators of diverse benthic invertebrates, providing strong top down control in marine coastal ecosystems and helping to maintain stability of canopy forming kelp forests across the Northeast Pacific Ocean (Arroyo-Esquivel et al., 2024; Galloway et al., 2023; Malakooti et al., 2026). Populations of *P. helianthoides* declined precipitously as the result of a *Vibrio pectenicida* FHCF-3 driven Sea Star Wasting Disease (SSWD) pandemic which co-occurred with the anomalously warm temperatures of the 2014 to 2016 Pacific Marine Heatwave (Aalto et al., 2020; Bond et al., 2015; Prentice et al., 2025). This led to functional extinction of *P. helianthoides* in the southern extent of their range from Baja Sur California, Mexico to Southern Oregon, USA releasing urchin herbivores from predation pressure, and leading to dramatic kelp forest deforestation, especially in Northern California (Arroyo-Esquivel et al., 2024; Galloway et al., 2023; Rogers-Bennett & Catton, 2019).

This dramatic population decline has resulted in *P. helianthoides* being listing as Critically Endangered on the IUCN Red List (Gravem et al., 2021) and Endangered by the Committee on the Status of Endangered Wildlife in Canada (COSEWIC, 2025). NOAA Fisheries proposed the species for listing as Threatened throughout its range (NOAA, 2023); however, to date, a final listing decision has not been made by the U.S. federal government.

In response to such a rapid and consequential population decline, a series of ongoing *P. helianthoides* research and conservation efforts have been identified, initiated, and fully implemented over the past decade (Heady et al., 2022). Importantly, recent work has led to the completion of the *P. helianthoides* life cycle in captivity, enabling for the formation of a captive breeding program (e.g. conservation aquaculture program) with dozens of aquaria actively rearing individuals (Hodin et al., 2021, 2025; Mizuta et al., 2023). These efforts have facilitated reintroduction experiments of captive reared individuals to understand the potential viability of future captive breeding programs. Furthermore, there are a variety of ongoing efforts to monitor and identify remnant intact populations across their known range, ranging from targeted population estimates via underwater census to community science driven programs like Snapshot Cal Coast (Rapacciuolo et al., 2021). Key to the success of management and recovery efforts are accurate inventories of remaining populations across their range. However, despite being a large and conspicuous benthic invertebrate, monitoring of this species over 12,000 km of coastline and up to 400 m of depth is not practical using visual methods (Lowry et al., 2022). Thus, new tools are needed to aid in ongoing monitoring efforts for *P. helianthoides*.

Over the past decade, targeted environmental DNA (eDNA) approaches have been demonstrated to be highly accurate, sensitive, and cost-effective tools for inventorying marine species (Beng & Corlett, 2020; Hunter et al., 2026; Langlois et al., 2021, 2025; Shelton et al., 2022; Taberlet et al., 2018). These approaches rely on the capture and identification of organism-specific nucleic acids, present as dissolved biomolecules or associated with particulate fractions in the water column (shed cellular debris or detritus) using sensitive and specific quantitative PCR (qPCR), droplet digital PCR (ddPCR), digital PCR, and CRISPR based approaches (Deiner et al., 2017; Williams et al., 2023). These molecular detection assays have limits of detection ranging from tens to hundreds of copies per liter of seawater. With this sensitivity, rigorously validated and thoughtfully implemented eDNA assays frequently outperform visual census approaches in detecting rare species per unit effort (Beng & Corlett, 2020; Fediajevaite et al., 2021; Iacaruso et al., 2025; Wikston et al., 2023). Furthermore, these approaches are highly cost-effective on the scale of $25 to $150 USD (in 2026) per sample and readily automatable with modern molecular facilities, enabling the scaling of these approaches for increased sampling efforts across time, space, and depth (Kelly et al., 2024). Increasingly, eDNA approaches are being utilized for quantitatively estimating biomass as a compliment to traditional surveys as there is frequently a predictable, low variability relationship between observed DNA copies and underlying biomass (Baetscher et al., 2025; Claver et al., 2026; Guri, Shelton, et al., 2024; Shelton et al., 2022, 2023). This is exemplified by the recent formal adoption of targeted eDNA approaches into Pacific Hake stock assessments (Hamel et al., 2025).

Sunflower sea stars have been previously detected by eDNA metabarcoding approaches utilizing primer sets that broadly target metazoan diversity (Jacquemot et al., 2024; Robinson et al., 2023; Shum et al., 2019). However, to date, no targeted assay has been developed for this species. Here we present *P. helianthoides-*specific qPCR and ddPCR assays that we fully validated across the five level validation scheme presented by Thalinger et al., 2021. Specifically, we conduct *in silico* and *in vitro* specificity and sensitivity analyses (Level 1); *in vitro* testing on closely related nontarget species (Level 2); detection obtained from environmental samples (Level 3); extensive field testing of environmental samples (Level 4); and comprehensive specificity testing, detection probability estimation from statistical modeling, and understanding ecological and physical factors influencing eDNA in the environment (Level 5). Furthermore, we present an externally valid framework building off of recent advances in eDNA assay design (Allison et al., 2023; Andruszkiewicz et al., 2020; Klymus et al., 2020; Lopez et al., 2025) to develop targeted eDNA assays for critically important species. Lastly, we provide detailed standard operating procedures for running qPCR and ddPCR single species assays adhering to Findable, Accessible, Interoperable, and Reproducible eDNA (FAIRe) standards (Takahashi et al., 2025) and Better Biomolecular Ocean Practices (BeBOP) guidelines (Pitz et al., 2026). These detailed, standards-driven protocols provide a valuable tool for enabling transparent reporting of eDNA data and results while adhering to FAIRe, Environmental Microbiology Minimum Information (EMMI), Darwin Core, and Minimum Information about any (X) Sequence (MiXS) standards to enhance biodiversity data mobilization efforts globally (Abarenkov et al., 2023; Borchardt et al., 2021; Klymus et al., 2024; Samuel et al., 2021).

## Methods

We present our *Pycnopodia helianthoides* assay design and methods in the context of an externally valid framework to develop targeted eDNA assays for critically important species (Table 1). First, we conducted a gap analysis of target and congener species across the known range of *Pycnopodia helianthoides* and organized existing sequencing data. Next, we filled in gaps through mitochondrial genome (mitogenome) skimming of missing species. We then designed the putative assay and validated it in the laboratory, mesocosms, and field.

**Table 1.** Overview of eDNA Assay Design.

| **Assay Design Steps** | **Steps Followed in This Current Project** |
| --- | --- |
| 1. Assemble Existing data | Organized existing assembled mitogenomes from National Center for Biotechnology Information (NCBI) GenBank. (n=114 Asteroidea mitogenomes)<br>Assembled existing whole genome sequencing (WGS) data from National Center for Biotechnology Information (NCBI) sequence read archive (SRA) (n=6 Asteroidea WGS archived datasets) |
| 2. *De novo* Sequence DNA Gaps | Conducted mitogenome skimming of missing target and related species across the *P. helianthoides*' range (n=20 mitogenomes, 13 taxa) |
| 3. Assemble Novel Mitogenomes for target and congeners | Assembled novel mitogenomes from 7 target and 20 non-target specimens |
| 4. Design Putative Assay | Employed *unikseq* to identify longest unique target region and Klymus et al. 2020 and Andruszkiewicz Allan et al. 2021 assay design in *Geneious Prime* |
| 5. Test on tissues | Confirmed assay specificity on 7 sea star species |
| 6. Determine Limits of Detection, Quantification and Blank | Systematically evaluated qPCR and ddPCR performance with synthetic DNA serial dilutions |
| 7. Test in mesocosm and field | Sampled University of Washington Friday Harbor Laboratories sunflower sea star holding tank and reintroduction experiment |
| 8. Paired with visual surveys | Paired Hakai Institute British Columbia Dive and eDNA Surveys |

### Assemble Existing data

We first assembled a list of described sea star species in the Northeast Pacific and Alaska (n=103 taxa, Table S1). We then screened NCBI GenBank for relevant mitogenome sequences for each of these individual species as well as each genera to identify sequencing gaps. All available Asteroidea mitochondrial genomes were downloaded (n=114 public Asteroidea mitogenomes), as well as available raw sequence reads from WGS projects so that mitochondrial genome assemblies from these data could be attempted (n=6 Asteroidea datasets).

### Sequence DNA Gaps

Our goal was to obtain mitogenomes from at least one of every co-occurring closely related genus to *P. helianthoides* based on the most recent phylogenetic understanding of Asteroidea to ensure adequate coverage of non-target co-occurring species (Jossart et al., 2024; Langlois et al., 2021; Mah & Blake, 2012; Sun et al., 2022). In order to fill in these apparent reference sequence gaps, we obtained tissue vouchers, genomic DNA, or existing sequence data from University of California Merced, Hakai Institute, and the California Academy of Science for 13 species (Table S2). Nucleic-acids were extracted from tissue samples using the Qiagen Blood & Tissue Extraction Kit. Genomic DNA for specimens without existing data was sequenced by Azenta Life Sciences using their standard non-human whole genome sequencing package on an Illumina NovaSeq X Plus with dual indices and paired 2x150 cycle chemistry, targeting 15 gigabases per specimen.

### Assemble Mitogenomes for target and congeners

Raw reads from novel sequencing and Illumina reads downloaded from NCBI SRA were quality controlled prior to mitogenome assembly. Raw read pairs were adaptor-trimmed of TruSeq3 adaptors using *Trimmomatic* v0.39 followed by merging using *FLASH* v1.2.11 (Bolger et al., 2014; Magoč & Salzberg, 2011). *Trimmomatic* was then used for quality score filtering using a 4 bp Q30 sliding window (minimum read length 50 bp) for all reads (merged, unmerged pairs, and unpaired reads after initial trimming). Quality-controlled merged and unmerged read pairs were used in subsequent mitochondrial genome assembly steps.

No single mitogenome assembly approach produced the best outcome in all circumstances. For this reason, the best mitogenome assembly was chosen from one of three (Illumina data) or one of two (PacBio data) approaches (choice indicated in Table S2). For Illumina reads: 1) quality-controlled reads were assembled with SPAdes v3.15.5 (-k 21,55,85,115,127; --isolate) (Prjibelski et al., 2020) and GetOrganelle v1.7.5 (get_organelle_from_assembly; -F animal_mt; --max-slim-extending-len 20000; --disentangle-time-limit 40000) (Jin et al., 2020) pulled the mitogenome from the assembly. 2) mitogenomes were assembled from quality-controlled reads within GetOrganelle v1.7.5 (get_organelle_from_reads; -F animal_mt; -R 30; --max-reads inf; --disentangle-time-limit 200000000; --reduce-reads-for-coverage inf) with no additional seeds outside of GetOrganelle’s default. 3) mitogenomes were assembled the same as #2, with the result of #2 provided as a seed for read recruitment. For PacBio reads (public data only): 1) mitogenomes were assembled using the same method as for the Illumina reads in approach #3. 2) PacBio HiFi reads were assembled using MitoHiFi v2 (-o 9; congener reference) (Uliano-Silva et al., 2023)The decision on the best approach was dependent on: 1) circularity and length of taxonomically accurate mitogenome, 2) number of final contigs, 3) convolution of the assembly path.

The chosen mitogenomes were annotated using the MITOS2 (Donath et al., 2019) web server on Galaxy (https://usegalaxy.org/). Half of the mitogenomes were missing an annotation for *trnF* from MITOS2, necessitating additional annotation using MitoZ (Meng et al., 2019). Gene boundaries were further curated through alignment with existing published mitogenomes from the Asteroidea using Geneious Prime v.2025 (Kearse et al., 2012). Mitogenome annotation visuals were generated in Geneious Prime.

The nuclear ribosomal RNA-internal transcribed spacer (rRNA-ITS) region was pulled from the SPAdes assemblies using Asteroidea 18S and 28S genes as bait sequences. Base discrepancies in multiple assembly paths were resolved through the use of ambiguous bases, resulting in one rRNA-ITS submission for each assembly when successful. These sequences are available on NCBI’s GenBank (see Table S2 for accessions).

To produce a high quality phylogenetic tree, we aligned the nucleotide sequences of 13 protein-encoding genes and 2 ribosomal RNA genes independently using MUSCLE 5.1 (Edgar, 2004) in Geneious Prime from 107 mitogenomes (8 *P. helianthoides*, 90 Asteroidea, 9 Echinodermata outgroups). Alignments were then masked to remove any position containing more than 30% gaps across the alignment and then concatenated (11,691 bp alignment). We then constructed a maximum likelihood phylogenetic tree using the GTRGAMMA model in *RAxML* 8.2.11 (Stamatakis, 2014), with 300 bootstrap replicates. The final tree was visualized using Iroki (Moore et al., 2020).

### Design Putative Assay

We used *unikseq* to identify unique, conserved gene regions to *P. helianthoides* (Allison et al., 2023; Lopez et al., 2025). Here we used the default parameters using 8 *P. helianthoides* mitogenomes as the target ingroup and for the outgroup using: 132 sea star mitogenomes (92 unique taxa, 80 species); 30 mitogenomes from commonly co-occurring and abundant echinoderm congeners; and 2 outgroups (Table S3). We then chose the longest identified unique fragment to *P. helianthoides* as the locus to design the hydrolysis probe based PCR assay. We followed recommended assay design guidelines of Andruszkiewicz et al., 2020 and Klymus et al., 2020 (Table S4). To understand the specificity of the designed assay, we isolated the primer and probe regions from a MUSCLE alignment of 135 *P. helianthoides* and Asteroidea *nad5* genes (Table S3), compared the frequency of mismatches, and used ggseqlogo v0.1 in R (Wagih, 2017) to calculate sequence conservation.

Next, we designed a synthetic positive control (gBlock) for the assay. The sequence included the PCR assay target with an appended short CO1 sequence from the extinct *Ectopistes migratorius* passenger pigeon to the ′5 end to enable contamination tracking (Tables S5,S6).

To test assay sensitivity *in vitro* <u>using quantitative PCR (qPCR),</u> we ran the assay with 8 replicates at each of the following concentrations of the synthetic gBlock (IDT) in copies per reaction: 200,000, 20,000, 2,000, 200, 20, 10, 5, 2. We note that we relied on the vendor supplied concentrations of purchased gBlock stocks (10 ng/µL) to generate all serial dilutions. We independently confirmed the concentrations of gBlocks via QuBit High Sensitivity Assay (ThermoFisher Scientific, USA). All measured concentrations were within .5 ng/µL of vendor supplied concentrations and thus assumed to be accurate (He et al., 2018). We used the recommended thermocycling conditions and reaction chemistry from the Taqman environmental master mix handbook (Tables 2-4). All qPCR assays were multiplexed using an internal positive control of *Euryapteryx curtus* broad-billed moa following Ramón-Laca et al., 2021 (Tables S6, S7). qPCR assays were run on an Applied Biosystem QuantStudio 5 at NOAA PMEL (Gold & Weinrich, 2026).

**Table 2.**
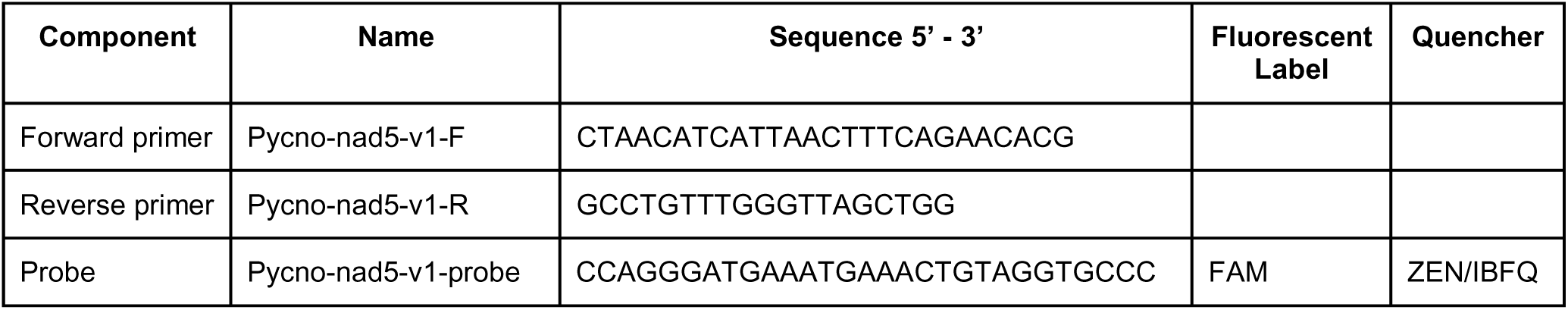
The designed *Pycnopodia helianthoides* hydrolysis probe PCR assay including sequences. Amplicon length is 123 bp.

**Table 3.**
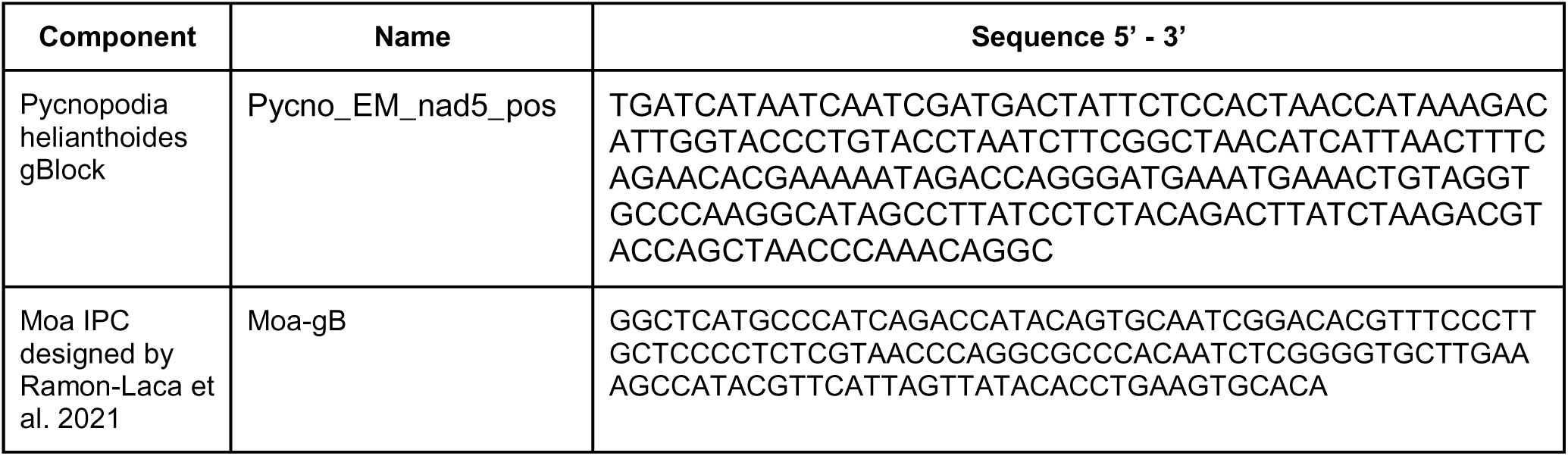
The designed *Pycnopodia helianthoides* and Moa Internal Positive Control synthetic gBlock sequences.

**Table 3.**
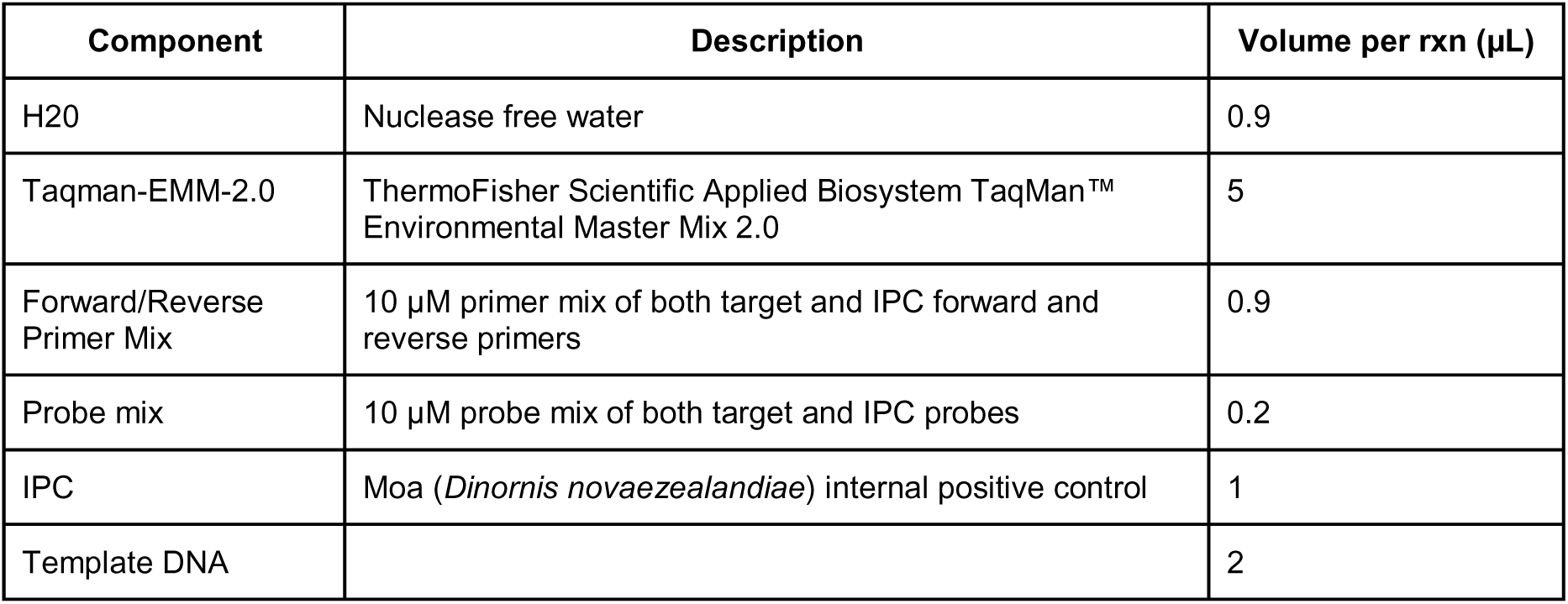
The *Pycnopodia helianthoides* qPCR assay reaction chemistry.

| Component | Description | Volume per rxn (μL) |
| --- | --- | --- |
| H2O | Nuclease free water | 0.9 |
| Taqman-EMM-2.0 | ThermoFisher Scientific Applied Biosystem TaqMan™ Environmental Master Mix 2.0 | 5 |
| Forward/Reverse Primer Mix | 10 μM primer mix of both target and IPC forward and reverse primers | 0.9 |
| Probe mix | 10 μM probe mix of both target and IPC probes | 0.2 |
| IPC | Moa (*Dinornis novaezealandiae*) internal positive control | 1 |
| Template DNA |  | 2 |

**Table 4.** The *Pycnopodia helianthoides* qPCR assay thermocycling conditions.

| Step | Name | Temperature (°C) | Time (s) | Cycles |
| --- | --- | --- | --- | --- |
| 1 | Initial Denaturation | 95 | 600 | 1 |
| 2 | Denaturation | 95 | 15 | 45 |
| 3 | Annealing | 60 | 60 |  |
| 4 | Hold | 10 | ∞ | 1 |

We also confirmed the performance of the primers and probe on Bio-Rad QX200 ddPCR Systems at Stanford University (Boehm Lab), California and at the Hakai Institute, British Columbia. Synthetic gBlocks at concentrations of 10 and 50 copies per µL were used during optimization testing of PCR conditions until the final method conditions (Table 5-6) yielded consistent expected results for the gBlocks (Gold & Weinrich, 2026).

**Table 5.**
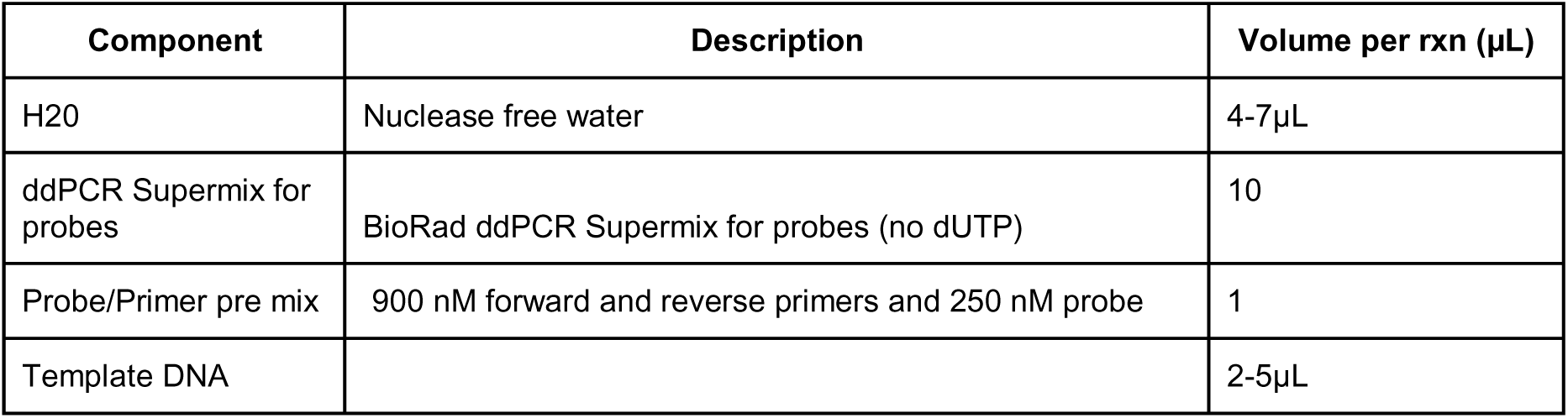
The *Pycnopodia helianthoides* ddPCR assay reaction chemistry.

| Component | Description | Volume per rxn (μL) |
| --- | --- | --- |
| H2O | Nuclease free water | 4-7μL |
| ddPCR Supermix for probes | BioRad ddPCR Supermix for probes (no dUTP) | 10 |
| Probe/Primer pre mix | 900 nM forward and reverse primers and 250 nM probe | 1 |
| Template DNA |  | 2-5μL |

**Table 6.** The *Pycnopodia helianthoides* ddPCR assay thermocycling conditions.

| Step | Name | Temperature (°C) | Time (s) | Cycle # | Cycles |
| --- | --- | --- | --- | --- | --- |
| 1 | Initial Denaturation | 95 | 600 | 1 | 1 |
| 2 | Denaturation | 95 | 30 | 2 | 35 |
| 3 | Annealing | 57 | 60 |  |  |
| 4 | Extension | 98 | 600 | 3 | 1 |

### Test on tissues

To confirm assay specificity relative to other sea stars expected to co-occur with *P. helianthoides* in the wild, we tested *Rathbunaster californicus*, *Leptasterias aequalis*, *Henricia pumila*, *Pisaster ochraceus*, *Patiria miniata*, and *Stylasterias ferrari* for cross amplification (Supplemental Materials 1) as well the target *P. helianthoides.* DNA or tube feet were obtained from UC Merced and Hakai Institute (Table S5). Nucleic acids were extracted from a single sample of each tissue using the same approach as described above for *P. helianthoides* and run as templates in both qPCR and ddPCR assays. We analyze 1:10 dilution, 1:100 dilutions, and 1:1000 dilutions of tissue derived extracted DNA extracts as template (2 µL) in qPCR assays (Table S8). Input nucleic-acids ranged from 0.12 to 18.8 ng template added per reaction. Reaction followed the protocol outlined in Tables 2-4.

### Determine Limit of Detection and Limit of Quantification

We conducted a series of serial dilutions to test for assay sensitivity for both the qPCR and ddPCR assays.

For the qPCR assay, we ran a serial dilution of the synthetic gBlock on three separate plates. The first plate contained a serial dilution of the following concentrations in copies per reaction: 200,000, 20,000, 2,000, 200, 20, 10, and 2. The plate also contained mesocosm and field samples from Friday Harbor Laboratories as described below. We observed complications with this initial serial dilution with high variance across replicates. This was the first time ever running a qPCR reaction in the NOAA PMEL Ocean Molecular Ecology laboratory - We then ran this plate a second time including the above described standards, mesocosm and field samples as well as tissues of *P. helianthoides* and other stars as described above. However, to improve accuracy of the LoB, LoD, and LoQ calculations we ran the assay on 24 replicates of a serial dilution of the synthetic gBlock (IDT) of the following concentrations in copies per reaction: 200,000, 20,000, 2,000, 200, 20, 10, 5, 2, 1.6, and 0.8. This third plate included the mesocosm and field samples from Friday Harbor Laboratories. For all standard curves, DNA copies were estimated from independent confirmation of input concentrations, not the vendor supplied concentration of 10 ng/µL. Specifically we used the QuBit fluorometry high sensitivity assay (ThermoFisher Scientific) to measure the concentrations of the 1:100 dilutions of vendor supplied concentrations. Observed concentrations were within 3 ng / µL expected vendor supplied concentration of 10 ng/µL and therefore we used our laboratory measured concentrations to generate standard curves (Table S9).

For the ddPCR assay, we similarly ran serial dilutions of the synthetic gBlock to calculate the LoD and LoQ at both the Hakai and Stanford labs. At Hakai, using 2 µL of DNA template, we ran the assay using the finalized ddPCR conditions (see Results), running five replicates of each of the following gBlock concentrations in copies per reaction: 625,000, 62,500, 12,500, 2,500, 500, 100, 20, 5, and 0.8, as well as an additional low-concentration curve spanning 2, 4, 6, 8, 10, 20, 40 copies per reaction. 625,000 and 62,500 gBlock standards saturated the assay, with positive droplets representing 93-100% of the total droplet count. At Stanford University, using 5 uL of DNA template, we ran the assay using the finalized ddPCR conditions (see Results), running eight replicates of each of the following gBlock concentrations in copies per reaction: 50,000, 5,000, 500, 50, 25, 5, 4, 2, and 1.

We calculated the Limit of Detection (LoD), Limit of Quantification (LoQ), and Limit of Blank (LoB), following both Lesperance et al., 2021 and Klymus et al., 2020 for qPCR and ddPCR approaches. Calculations for ddPCR limits following Klymus et al. 2020 used the estimated concentrations derived from the instrument whereas limits following Lesperance et al. 2021 using the number of positive and total analyzed droplets given the reliance on binary data for eLowQuant calculations.

### Test in mesocosm & field

On May 16th, 2025, we tested the qPCR assay on eDNA samples collected from the common outflow of the Friday Harbor Laboratories’ sunflower sea star flow through system (Table S10). The flow through system has a flow rate of approximately 0.85 liters per second. At the time of sampling, the aquarium contained 39 adult *P. helianthoides* stars larger than 40 cm in diameter. We collected eDNA samples using peristaltic pressure filtration of one liter using a Smith Root PES 47 mm diameter 0.22 µm pore size filter (https://doi.org/10.5281/zenodo.17655098). We compared this approach against different active and passive filter approaches (see supplemental materials).

On May 16th, 2025, we collected three replicate one liter surface seawater samples adjacent to a location where captive reared *P. helianthoides* had been reintroduced in summer 2024, adjacent to Friday Harbor Laboratories’ pier (48.545 N, -123.012 W). We also collected triplicate one liter samples near the seawater intake pipe approximately 500m east of the pier, which we expected to be a negative or low dose control. However, a singular *P. helianthoides* juvenile individual was observed at 10 m depth near this location within one week after eDNA sample collection. Samples were collected in bleach washed, deionized water rinsed, sterilized plastic Nalgene bottles that were triple rinsed with ambient seawater prior to sample collection. Samples were then filtered through a Sterivex filter to collect eDNA, and preserved, as above.

Nucleic-acids were extracted from all filters obtained from mesocosm and field eDNA samples at NOAA PMEL using a modified Qiagen Blood and Tissue Kit (sterivex extraction protocol:https://zenodo.org/records/17655148). Sterivex filters were extracted on May 22nd, 2025.

Mesocosm and field nucleic-acid extracts were run as templates in four replicate qPCR assays on the LoD and LoQ plate using the protocol described above. We included 6 no template controls (NTC) consisting of molecular grade, nuclease free water. All samples, NTCs, and standards were run with the internal positive control described above.

Output values from the qPCR machine were then converted to copies per L for the original water sampled using the following equation (Equation 1):

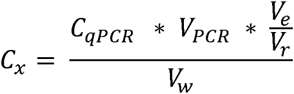

Where C_qPCR_ is the concentration generated by the qPCR machine (in copies per µL), V_PCR_ is the volume of the total PCR reaction (in µL) here 10 µL. V_e_ is the volume post DNA extraction (in µL) here 100 µL. V_r_ is the volume of template added to the PCR reaction (in µL) here 2 µL. V_w_ is the volume of sample water filtered onto the sterivex for extraction (in liters).

### Paired with visual surveys

Field testing and comparison with visual surveys was carried out using eighty 1-liter seawater samples containing potentially endogenous, native sunflower sea stars that were filtered on 0.22 µm pore size Sterivex filters following the protocol described in Kellogg, 2024 using Longmire’s buffer to preserve the DNA until extraction. Sixty of these samples were collected from subtidal regions of Calvert Island, British Columbia with paired dive surveys that recorded the number and estimated biomass of *P. helianthoides* (see dive methods in Gehman et al., 2025). Biomass density at the transect level was calculated as kg per 10 m^2^ using a length–biomass conversion from (Lee et al., 2016). Of those 60 samples, 34 were from fjord locations while 26 were from the outer islands around Calvert Island. In addition, 20 intertidal samples were selected for inclusion because we suspected there to be no sea stars present given that *P. helianthoides* has become rare in the intertidal regions. Twelve intertidal samples were from outer Calvert Island sites, while eight were from Quadra Island. All intertidal samples have paired transect survey data.

Sites were divided into four groups (referred to as regions): subtidal surveys in fjords by Calvert (Subtidal - Fjords), subtidal surveys around outer islands by Calvert (Outer Islands - Subtidal), intertidal samples from the outer island of Calvert (Outer Islands - Intertidal) and intertidal samples from Quadra (Quadra - Intertidal, Figure 1). These data were collected on the traditional territories of the Wuikinuxv Nation, the Heiltsuk Nation, the Nuxalk Nation and the We Wai Kai Nation, who hold indigenous rights to their territories.

**Figure 1.**
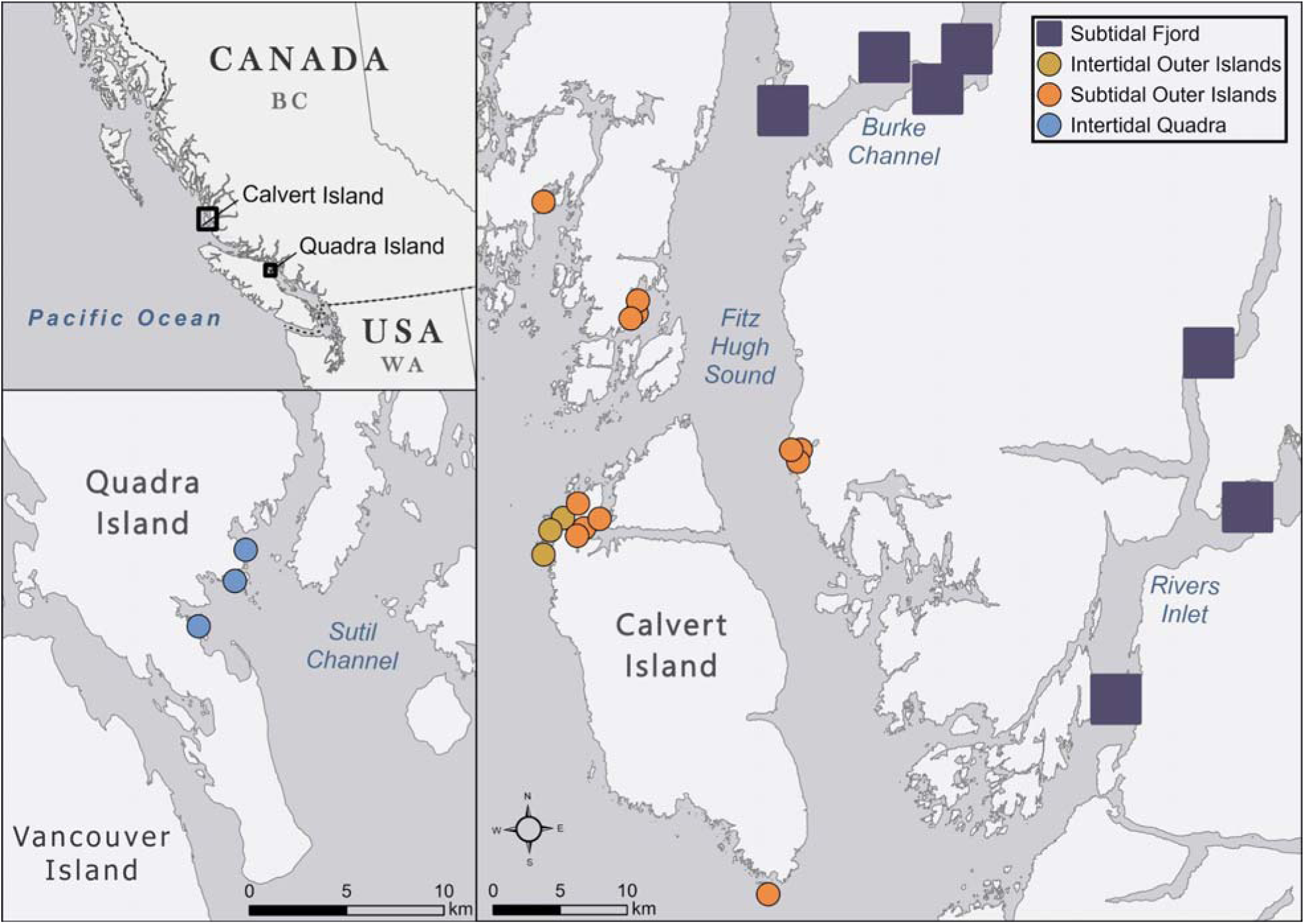
Map of paired visual - eDNA survey study area. Upper left depicts the regional map with Quadra Island inset in lower left and Calvert Island inset to the right. Locations are divided into four subgroups: Subtidal Fjord, Subtidal Outer Islands, and Intertidal Outer Islands of Calvert Island and Intertidal at Quadra Island. Note that the fjord sites are part of a data sharing agreement with local First Nations, and coordinates are only given to a 4 km resolution (blue square).

DNA was extracted using ZymoBIOMICS DNA miniprep kits in clean rooms designated for the processing of environmental DNA at Hakai Institute’s Genomics Laboratory. To adapt this kit for use with Sterivex filter units, BashingBeads™were added directly to the filter unit, along with 750 µl ZymoBIOMICS Lysis Solution (Zymo Research, USA) and vortexed for 20 minutes. Lysate (and beads) were removed from the Sterivex into a 2mL tube by centrifuging with the Sterivex inlet into a 2 mL tube (i.e. upside down) placed inside a 50mL conical tube at 3000 x g for 3 mins and then the extraction proceeded as per manufacturer’s instructions. Droplet digital PCR (ddPCR) was carried out using three replicates of each DNA sample along with two no-template controls and two replicates each of synthetic gBlock standards at concentrations of 250 copies per uL, 50 copies per uL and 10 copies per uL as positive controls. Any samples that had less than 12,000 acceptable droplets were labeled as failures and re-run. Due to high initial concentrations for some of the samples, a subset of samples were diluted 10-fold and 50-fold and rerun to obtain a more accurate estimate of *P. helianthoides* DNA concentration. Output values from the ddPCR machine were then converted to copies per L for the original water sampled using the following equation (Equation 2):

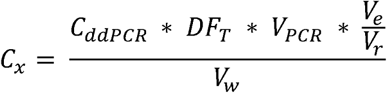

Where C_ddPCR_ is the concentration generated by the ddPCR droplet reader (in copies per µL), DF_r_ is the dilution factor, if any, that the extracted DNA was diluted by before adding to the PCR reaction. V_PCR_ is the volume of the total PCR reaction (in µL). V_e_ is the volume post DNA extraction (in µL). V_r_ is the volume of template added to the PCR reaction (in µL). V_w_ is the volume of sample water filtered onto the sterivex for extraction (in liters).

### Opportunistic Wasting versus Healthy Comparison

Between 2018 and 2021, no sunflower sea star observations were reported across California, USA. However, beginning in 2022, sunflower sea stars were observed north of San Francisco Bay, first in Humboldt, Mendocino and most recently in Sonoma counties (MARINe.org, iNaturalist, social media, and personal communications). On August 17, 2025, opportunistic eDNA samples were collected from an aggregation of individuals in Sonoma County, CA. Seawater samples were collected approximately 1 m above one *P. helianthoides* individual without discernable signs of SSWD and one *P. helianthoides* adult individual with clear characteristics of SSWD (e.g. multiple arm loss, open lesions on aboral surface). Additional eDNA samples were collected 50 m from the wasting individual as well as 50 and 100 m from the healthy individual. Seawater samples were collected on SCUBA using single-use 1-liter enteral feeding pouches (Covidien, Dublin, Ireland). On shore, samples were gravity filtrated onto 0.22 µm pore size PVDF Sterivex cartridges following the methods of (Shea & Boehm, 2023b). The specific amount of water filtered varied by sample (between 400-500 mL) and was recorded (Table S11). Filters were immediately frozen at -20°C until DNA extractions, which occurred within 5 months of sample collection.

Nucleic-acids were extracted from all filters from these opportunistic samples at Stanford University using the Qiagen DNeasy Blood and Tissue Kit (Qiagen, Germantown, MD, USA) with modifications (Shea & Boehm, 2023a; Spens et al., 2017). Nucleic-acid extracts were run as template (5 µl) in three replicate ddPCR reactions using the finalized ddPCR conditions (see Results) and an AutoDG Automated Droplet Generator (Bio-Rad, Hercules, CA), a C1000 Touch Thermal Cycler (Bio-Rad, Hercules, CA) and a QX200 Droplet Digital PCR System (Bio-Rad, Hercules, CA) (Shea et al., 2026). We included 1 no-template control (NTC) consisting of molecular grade, nuclease free water and 1 positive control consisting of a synthetic gBlock (IDT) at 50 copies per µL template. Output values from the ddPCR machine were then converted to copies per L for the original water sampled using Equation 2.

### Opportunistic Larval eDNA Generation

On January 23, 2026, opportunistic water samples were collected to characterize larval *P. helianthoides* shedding dynamics. In advance, we created 6 mesocosms containing 10 liters of sterilized sea water with 10,000 larvae, 1,000 larvae, 100 larvae, 10 larvae, 1 larva, and no larvae (mesocosm blank). The larvae used in these samples were mid-late stage bipinnaria larvae 16 days post fertilization cultured at 16-17°C. All the larvae used for this sample collection came from the same original culture vessel (22 L carboy) at the Sunflower Star Laboratory. To generate sterile sea water for mesocosms, natural seawater from an offshore intake in Monterey Bay, CA was 0.5 µm pore size filtered and UV sterilized. Larvae were fed a mono algal diet of *Rhodomonas salina* with daily water changes via forward filtration. Mesocosms were prepared 24 hours prior to sample collection and larvae were stocked in static tubs with no aeration and fed *R. salina* at approximately 15 cells / µL seawater. We then collected three replicate 1-liter water samples from each mesocosm directly into single-use enteral feeding pouches (Covidien, Dublin, Ireland) by siphoning water through a 100 µm mesh to exclude larval bodies from the samples; mesocosms were stirred while siphoning to promote even distribution of shed DNA. We prepared 6 sets of tubing, mesh, and stirring sticks that were all sterilized in 10% bleach solution and rinsed in sodium thiosulphate solution prior to sampling. Filled enteral feeding pouches were then used for gravity filtration onto 0.22 µm pore size, PVDF Sterivex cartridges, and filters were immediately stored at -20°C (Shea & Boehm, 2023b). We also directly pipetted 1 individual larva and 10 individual larvae onto 0.22 µm pore size, PVDF Sterivex cartridges in triplicate, to measure the DNA concentrations that might be generated if an entire larval body were to be captured in an eDNA sample; these filters were also immediately stored at -20°C.

Within 10 days of sample collection, nucleic-acids were extracted from all filters and run as template (5 µl) in two replicate ddPCR assays at Stanford University using the same methods described above for the healthy and wasting star comparison. Due to high initial concentrations in some of these larval samples, a subset of nucleic-acids were re-run at a 1:20 dilution as template; dilutions were done using nuclease-free water.

## Results

### Assembled Mitogenomes for Target and Congeners

We obtained 1.04 billion reads (min = 3.5 million, max = 281 million, mean = 52 million) across the 20 newly sequenced specimens (Table S2). We successfully generated 27 novel mitochondrial genomes, representing 19 sea star species, including 7 individuals from *Pycnopodia helianthoides* from across the species’ historic range (Figure 2, Figure S1). The assembled mitogenomes include 6 Asteriodea (plus 1 Echinodermata outgroup) assembled from public NCBI SRA records (Table S2). Coverage of generated mitogenomes ranged from 8x to 9,474x with a mean of 872x and GC content ranged from 26.9% to 38.9% with a mean of 35.0% (Table S2, Supplemental Figure S2). In addition to these 27 new mitogenomes, we also downloaded 114 existing Asteroidea and 31 outgroup mitogenomes from NCBI GenBank. Combined, the sea star mitogenome dataset encompasses 140 mitogenomes, 93 unique taxa, and 81 species), including 28 of the closest related co-occurring congeners in the sea star Family Asteriidae along the Northeast Pacific Coast. We assembled a phylogenetic tree from these mitogenomes showing *P. helianthoides* is monophyletic (Figure 3).

**Figure 2.**
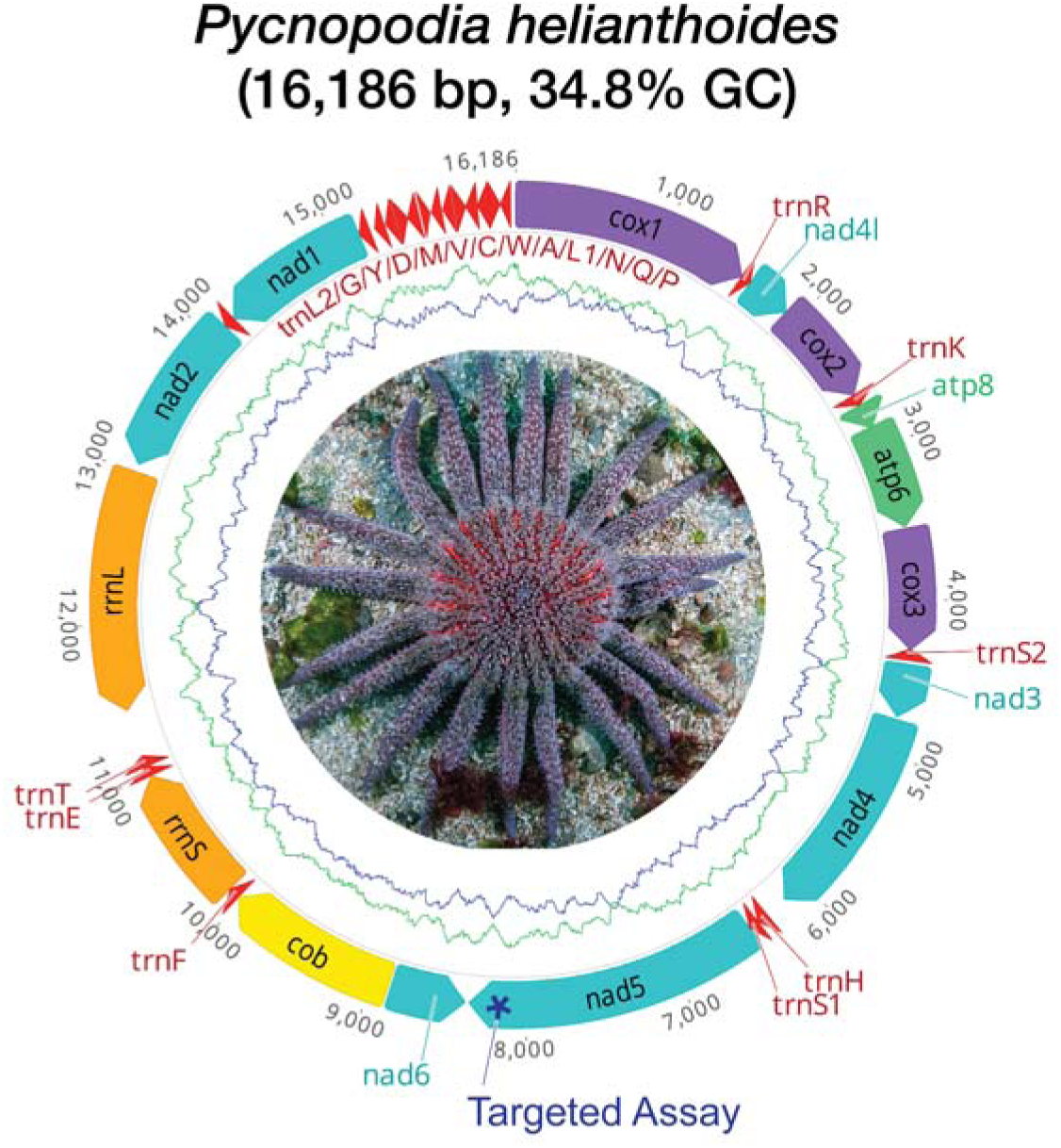
Assembled mitochondrial genome of *Pycnopodia helianthoides*. Circular mitogenome map of *Pycnopodia helianthoides* (specifically M0D062475I, accession XXX). All annotated genes are indicated with their direction. Inner tract shows GC content (green) and AT content (purple) with an 80□bp sliding window. The target assay is located near the terminal end of the *nad5* (ND5/NADH5) gene 8,162 bp to 8,284 bp on the mitogenome (123 bp amplicon).

**Figure 3.**
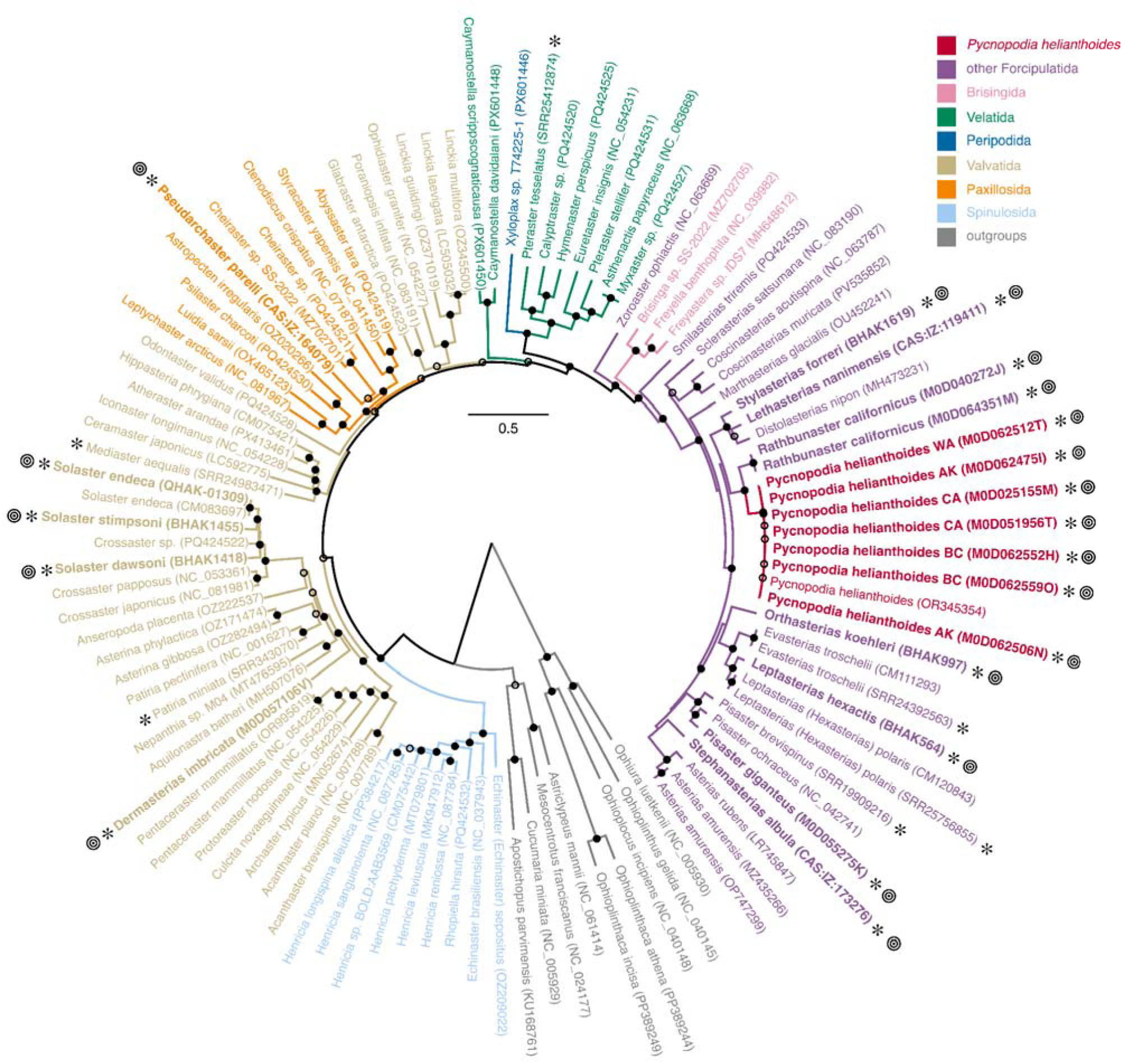
Mitogenomes of class Asteroidea. Multi-gene phylogenetic tree (n = 15 genes, 107 mitogenomes) showing the available reference sequences of Asteroidea. Colors indicate different Asteroidea orders, red for *Pycnopodia helianthoides,* and grey for other echinoderm taxa used as outgroups. Novel mitogenomes generated for this study are indicated with asterisks (*), with newly sequenced specimens also indicated in boldface with target symbols (□). Bootstrap support is indicated with open (>50%) and closed (>80%) dots at nodes.

### Assay Design

We identified 45 unique *P. helianthoides* sequence fragments larger than 100 bp using *unikseq* (max = 632 bp, mean 197 bp). The longest unique fragment mapped to the NADH dehydrogenase 5 (*nad5*) gene and was 100% unique to *P. helianthoides* across the sea star mitogenome database assembled here. In addition, this region was nearly identical across all 8 *P. helianthoides* individuals with the average proportion (%) of ingroup entries 99.7% of shared reference k-mers over unique region length. We then designed a primer pair and probe for a 123 bp fragment of this unique *nad5* gene region that met all recommended criteria outlined in Andruszkiewicz et al., 2020 and Klymus et al., 2020 (Figure 4, Figure S3, Table S4).

**Figure 4.**
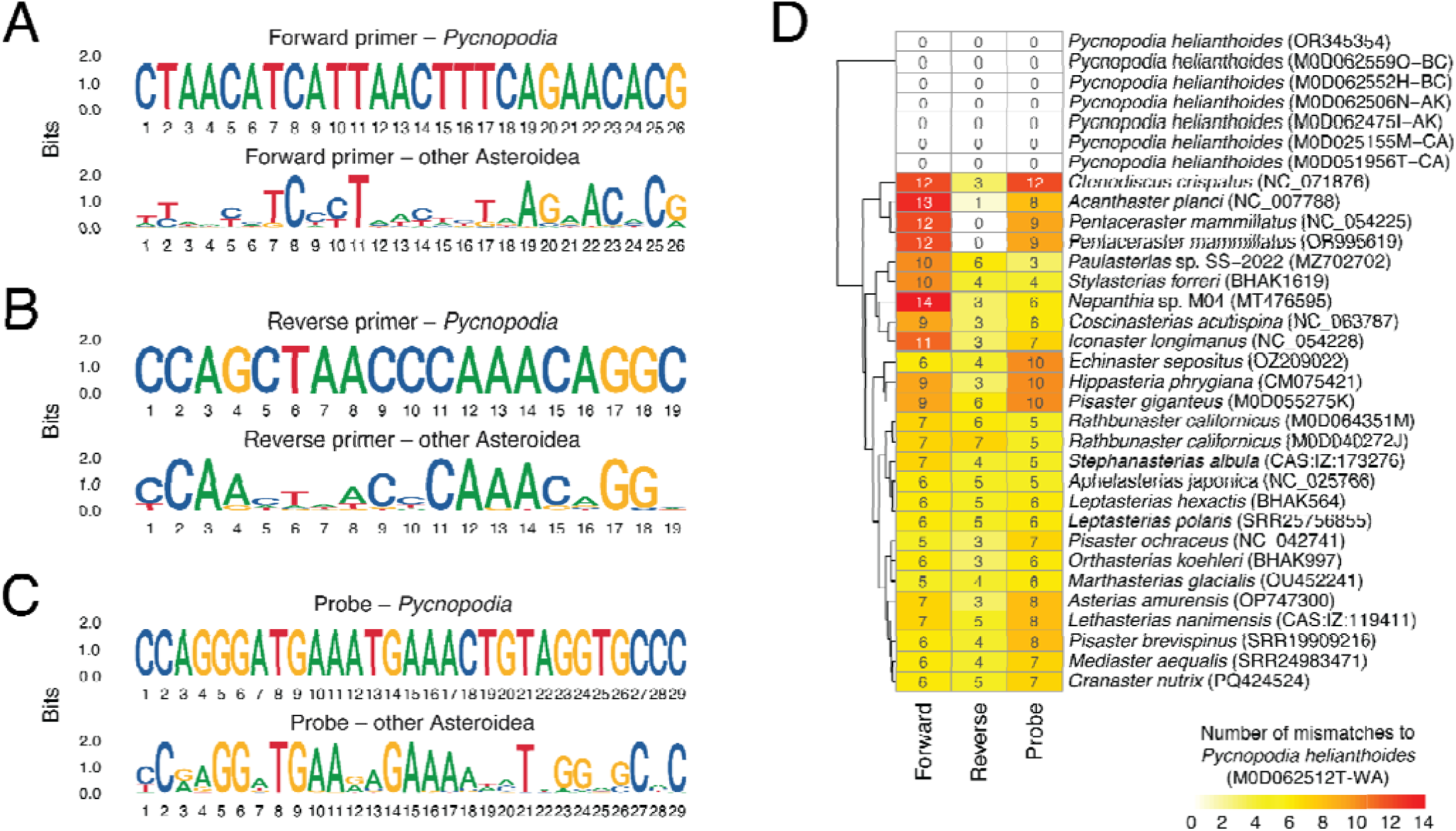
Assay Design Specificity. The *Pycnopodia helianthoides* assay is specific to the species, inclusive of all regional representatives, and exclusive of close and distant relatives. Sequence logos for forward primer (A), reverse primer (B), and probe (C) regions of the *P. helianthoides* qPCR assay comparing intra- and interspecific sequence conservation. The bit score is the informational entropy, with a higher bit score corresponding to a conserved position and a lower bit score corresponding with divergence. Counts of mismatches are compared (D) for the most closely related sea stars as well as species with the fewest mismatches for each primer and probe (Table S12). NCBI Accession numbers or voucher IDs provided in parentheses.

### Assay Validation

We validated the designed primer and probe for use as both qPCR (Tables 3 & 4) and ddPCR (Tables 5 & 6) using both synthetic oligos and nucleic-acids from *P. helianthoides* tissues.

### Test on tissues

We confirmed assay specificity *in vitro* observing a lack of cross amplification of co-occurring sea stars *Rathbunaster californicus*, *Leptasterias aequalis*, *Henricia pumila*, *Pisaster ochraceus*, *Patiria miniata*, and *Stylasterias ferrari* tissue derived DNA extracts observing no amplification after 40 cycles of qPCR across all species, dilutions, and all replicates (LoD = 0.2 Copies per reaction). This qPCR plate experiment had successful amplification of both *P. helianthoides* standards and internal positive controls within each tissue sample with no evidence of inhibition. All no template controls failed to amplify. We note that in the testing, one specimen of *P. miniata* amplified at low concentrations; an alternative tissue was sourced that did not amplify. This species had 10 mismatches each for forward primer and probe and 5 mismatches for the reverse primer. Given the high number of mismatches between the assay components and *P. miniata*, we assumed this was a sample specific cross contamination event.

### Assay LoD/LoQ

#### qPCR

The Limit of Blank was observed to be 0 copies per reaction across all negative controls. We calculated LoD and LoQ for each plate. Following Lesperance et al. 2021 (e.g. eLowQuant) method across 24 replicate standard curves, we determined the LoD of the qPCR assay was 0.2 copies per reaction and the LoQ was determined to be 0.8 copies per reaction. Following Klymus et al. 2020 method across 24 replicate standard curves, we determined the LoD of the qPCR assay to be 3.7 copies per reaction and the LoQ to be 150 copies per reaction. We note that there was considerable variation in LoD and LoQ per plate, particularly our first plate which displayed clear signs of pipetting error in a handful of wells of the internal positive control.

#### ddPCR

The Limit of Blank was observed to be 0 copies per reaction across all negative controls. Following Lesperance et al. 2021 method, we determined the LoD of the ddPCR assay was 0.1 copies per reaction at Stanford and 0.5 copies per reaction at Hakai accepting a single observed droplet constitutes a positive detection. The limit of detections were both about 6 times higher when requiring at least three observed droplets constitute a positive detection, a common practice in ddPCR analyses (Table 7). The LoQ was determined to be 0.5 copies per reaction at Stanford and 1.7 copies per reaction at Hakai. Following Klymus et al. 2020 method, we determined the LoD of the ddPCR assay to be 3.3 copies per reaction and the LoQ to be 11 copies per reaction at Stanford whereas at Hakai we observed an LoD of 6.7 copies per reaction and an LoQ of 16 copies per reaction when considering the low-range curve or an LoD of 2.1 copies per reaction and an LoQ of 16.0 copies per reaction with the full-range curve. We note that there was minimal variation in LoD and LoQ per plate.

**Table 7.** Estimated Limits of Blank, Detection, and Quantification for qPCR and ddPCR approaches. Limits were calculated following both Lesperance et al. 2021 and Klymus et al. 2020 with units in copies per reaction. qPCR and ddPCR had similar LoB, LoD, and LoQ within each method. However, there are significant differences in both LoD and LoQ between the Lesperance et al. 2021 and Klymus et al. 2020 methods. Hakai Low indicates the low-range gBlock curve (2-40 copies per reaction) while Hakai Full indicates the full-range gBlock curve (0.8-625,000 copies per reaction).

| Limit Method | Assay Type | Droplets | Lab | Plate | Template Volume | Reaction Volume | LOB | LOD | LOQ |
| --- | --- | --- | --- | --- | --- | --- | --- | --- | --- |
| eLowQuant | qPCR | - | NOAA PMEL OME | Pycno Run 5 | 2 | 10 | 0 | 0.2 | 0.8 |
| eLowQuant | ddPCR | 1 | Stanford | Stanford | 5 | 20 | 0 | 0.1 | 0.5 |
| eLowQuant | ddPCR | 2 | Stanford | Stanford | 5 | 20 | 0 | 0.2 | 0.7 |
| eLowQuant | ddPCR | 3 | Stanford | Stanford | 5 | 20 | 0 | 0.6 | 2.2 |
| eLowQuant | ddPCR | 1 | Hakai | Hakai Low | 2 | 20 | 0 | 0.5 | 1.7 |
| eLowQuant | ddPCR | 2 | Hakai | Hakai Low | 2 | 20 | 0 | 1.2 | 4.6 |
| eLowQuant | ddPCR | 3 | Hakai | Hakai Low | 2 | 20 | 0 | 2.9 | 11.1 |
| Klymus | qPCR | - | NOAA PMEL OME | Pycno Run 1 | 2 | 10 | 0 | 25.4 | 786.0 |
| Klymus | qPCR | - | NOAA PMEL OME | Pycno Run 3 | 2 | 10 | 0 | 6.0 | 41.0 |
| Klymus | qPCR | - | NOAA PMEL OME | Pycno Run 5 | 2 | 10 | 0 | 3.7 | 150.0 |
| Klymus | ddPCR | 1 | Stanford | Stanford | 5 | 20 | 0 | 3.3 | 11.0 |
| Klymus | ddPCR | 2 | Stanford | Stanford | 5 | 20 | 0 | 12.0 | 12.0 |
| Klymus | ddPCR | 3 | Stanford | Stanford | 5 | 20 | 0 | 9.7 | 24.0 |
| Klymus | ddPCR | 1 | Hakai | Hakai Low | 2 | 20 | 0 | 6.7 | 16.0 |
| Klymus | ddPCR | 2 | Hakai | Hakai Low | 2 | 20 | 0 | 10.0 | 17.0 |
| Klymus | ddPCR | 3 | Hakai | Hakai Low | 2 | 20 | 0 | 6.3 | 17.0 |
| Klymus | ddPCR | 1 | Hakai | Hakai Full | 2 | 20 | 0 | 2.1 | 16.0 |
| Klymus | ddPCR | 2 | Hakai | Hakai Full | 2 | 20 | 0 | 23.9 | 23.9 |
| Klymus | ddPCR | 3 | Hakai | Hakai Full | 2 | 20 | 0 | 19.5 | 19.5 |

### *Pycnopodia helianthoides* Successfully Detected in Mesocosm

We observed ∼2 million copies per L of Pycnopodia target DNA in the Friday Harbor Aquarium discharge water (mean ± sd = 2,330,548 ± 1,657,826 copies per L). The discharge is from a flow through system with 39 individual stars of at least 40 cm in diameter. Sterivex 0.22 µm filters yield the highest concentration of Pycnopodia DNA (Figure S4). All observations were multiple orders of magnitude above the LoD and LoQ (Table 1).

### *Pycnopodia helianthoides* Successfully Detected in field

We detected the eDNA in water samples collected adjacent to 10 reintroduced stars from at least one technical replicate from each of the three surface eDNA samples taken ∼5 m above the individuals on three replicate PCR plates (Figure 5). However, we note that these detections were never above the Klymus et al. 2020 calculated LoD, but were above the LoB, meaning that fewer than 95% of replicate samples at this concentration are likely to be amplified. In contrast, all of these observations were above the Lesperance et al. 2021 calculated LoD. We also found a higher number of detections on the earlier plates with 2 and 4 freeze thaws of the extracted DNA as compared to the plate using template DNA that experienced 5 freeze thaws. All of the detections from the second plate with four freeze thaws were after 40 cycles.

**Figure 5.**
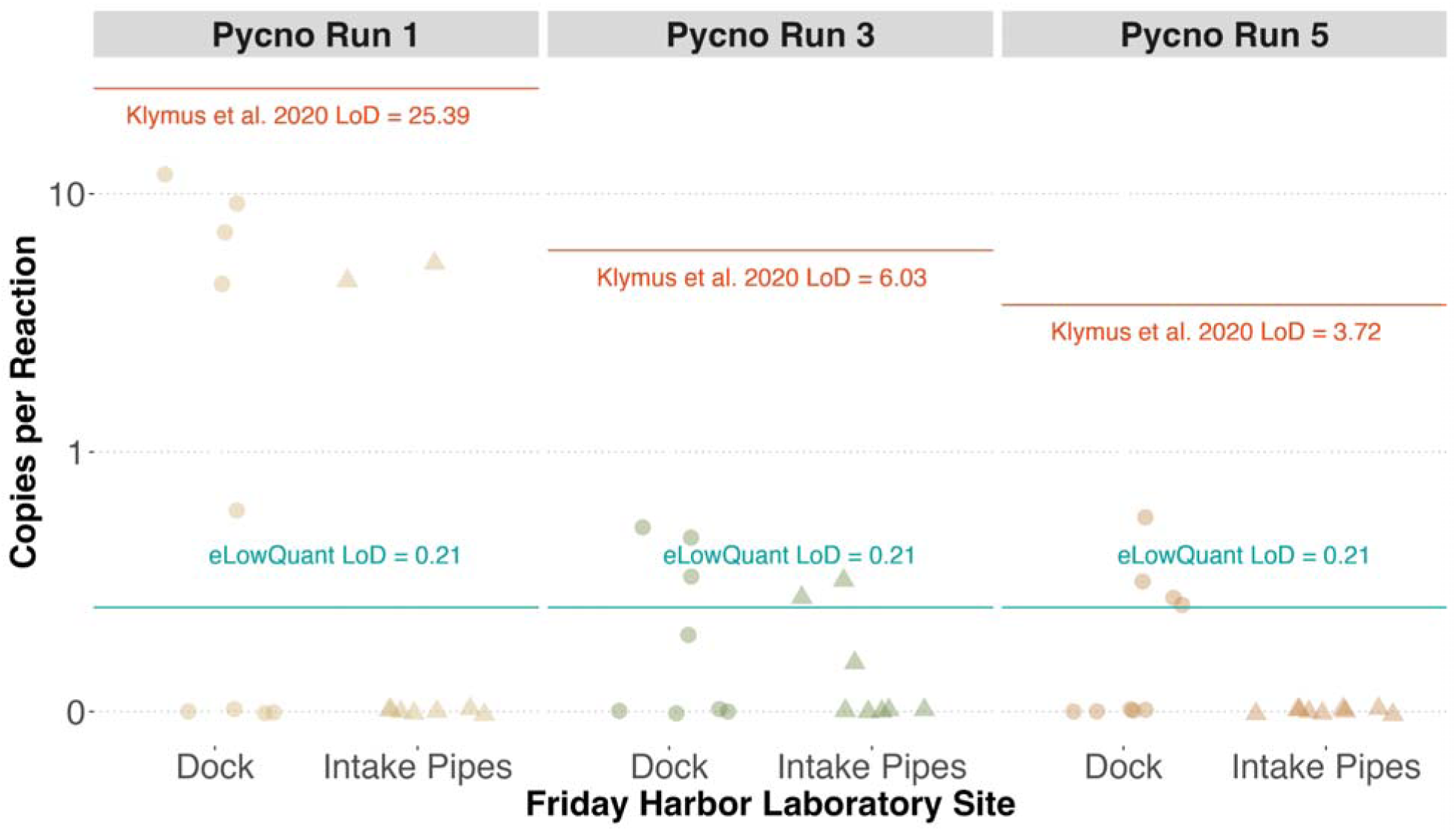
Detection of sunflower sea star eDNA from mesocosm and field experiments. Sunflower sea stars were detected from individuals reintroduced under the University of Washington Friday Harbor Lab (FHL) dock as well as the FHL intake pipes. Pycno Run 1 was loaded with template DNA that experienced two freeze thaws where Pycno Run 3 and Pycno Run 5 had experienced four and six freeze thaws respectively. Detections were above the Lesperance et al. 2021 eLowQuant limit of detection (LoD), but not the Klymus et al. 2020 LoD. All detections were above the limit of blank (0 copies per reaction). These results highlight the importance of the number of template freeze thaws and methods for calculating the limit of detection on the interpretation of eDNA signatures.

These results were also mirrored by observations at the Friday Harbor Lab intake pipes where only a single individual star was observed. Here we detected eDNA of *P. helianthoides* from one of three surface samples taken at the Friday Harbor Lab intake pipes only from the first PCR that was loaded with template DNA that only underwent two freeze thaw cycles (Figure 5). Similarly, these samples had a concentration below the Klymus et al. 2020 calculated LoD, but above the Lesperance et al. 2021 calculated LoD.

These results highlight the importance of both freeze thaw cycles and methodological choice of limit of detection calculations on ecological interpretation of low concentration eDNA.

### Opportunistic Wasting versus Healthy Comparison

Similar to the translocation observations at Friday Harbor Laboratories, where many samples were close to the LoD (meaning that the specific LoD used changed the interpretation of the results), most of the samples collected near healthy and wasting individual stars in Sonoma, CA fell between our most conservative and most liberal calculated LoDs. The samples collected 0 m, 50 m, and 100 m from the healthy individual had DNA concentrations ranging from 63 copies/L to 540 copies/L (average: 236 copies/L, SD: 134 copies/L). Similarly, the sample collected 50 m from the wasting individual was 129 copies/L (SD: 67 copies/L). The observed LoD for ddPCR results conducted at Stanford University ranged from 4 copies per L (eLowQuant method, single positive droplet, largest sample volume) to 600 copies per L (Klymus et al. 2020 method, two positive droplets, smallest sample volume). In contrast, the sample collected directly above the wasting individual had a DNA concentration of 52,524 copies/L (SD: 5853 copies/L), two orders of magnitude higher than all other field samples collected during this opportunistic sampling campaign and far above all calculated LoQs.

### Opportunistic Larval eDNA Generation

Across 10 liter mesocosms containing 1 larva to 10,000 larvae, we were able to successfully detect *P. helianthoides* eDNA (Figure 6). DNA concentrations detected in the mesocosms ranged from 9247 copies/L (SD: 971 copies/L) for 1 larva to 4,919,718 copies/L (SD: 1,250,284 copies/L) for 10,000 larvae), well above all calculated LoDs. When extracting DNA directly from sterivex filters containing larval bodies, we found significant variation in the copies of DNA generated from a single individual; triplicate samples containing 1 larva and 10 larvae ranged in concentration from 407,735 - 1,565,958 copies/individual (SD: 473,753 copies/individual).

**Figure 6.**
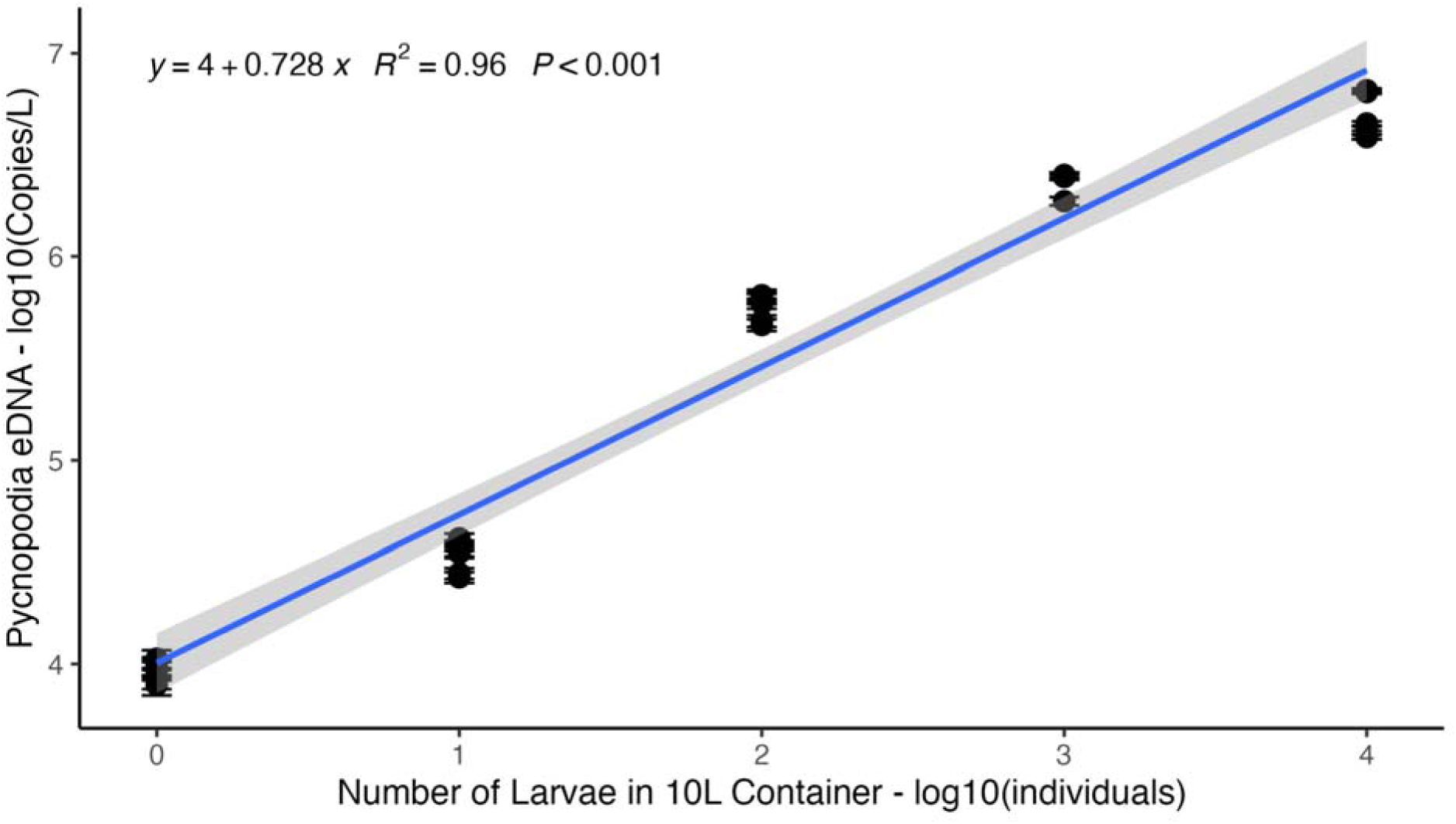
eDNA concentrations in larval mesocosm study. Concentration of eDNA is strongly correlated with the number of *Pycnopodia helianthoides* larvae in a sea water tank. These results highlight that *P. helianthoides* larvae may serve as a source of eDNA.

### Paired with visual surveys

Subtidal visual surveys observed the most *P. helianthoides* in the fjords near Calvert Island with only one site having zero individuals on transect. In contrast, the outer islands subtidal sites had *P. helianthoides* present at 73% of sites. Visual surveys only rarely observed *P. helianthoides* in intertidal sites with its presence only observed on one survey on Quadra Island (Figure 6).

The ddPCR eDNA assay detected *P. helianthoides* DNA at all subtidal fjord sites and 80.8% of subtidal outer island sites. For the intertidal sites, eDNA-ddPCR results showed 50% detection of *P. helianthoides* at Calvert Island intertidal sites and only 12.5% of Quadra Island intertidal sites (Figure 7). The range of eDNA concentrations ranged from below our limit of detection (450 copies/L) to above 800,000 copies/L resulting in log10 transformation of the data in order to visualize patterns. Several of the fjord samples were so high in concentration that a 10-fold dilution of the DNA template was necessary toto avoid oversaturation in the ddPCR assay.

**Figure 7.**
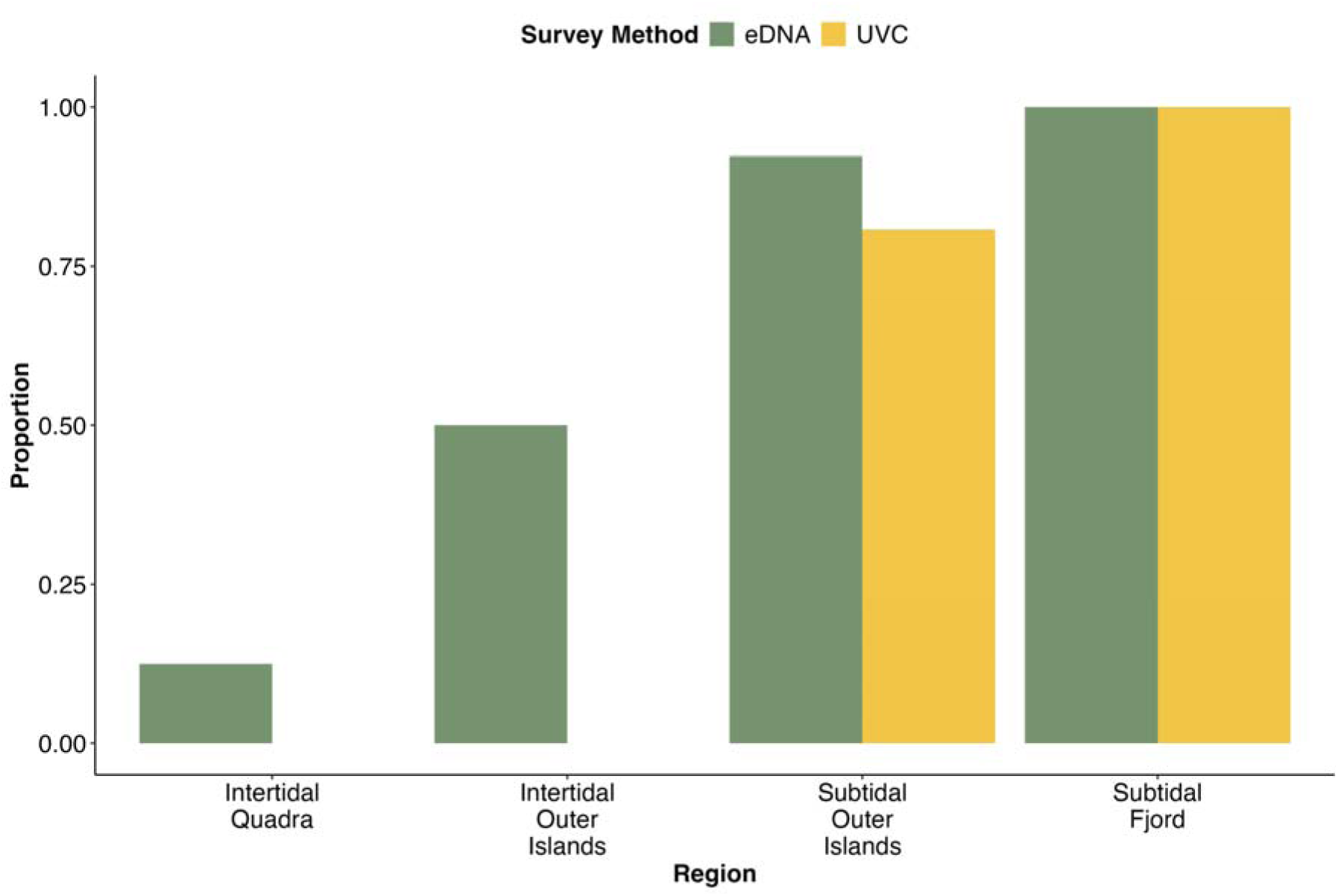
Proportion of Observations with Sunflower Sea Star Detections. Proportion of samples or dive transects across the four regions with a positive detection of *Pycnopodia* by either underwater visual census (UVC) or by ddPCR assay of water samples (eDNA). Samples below LoD were set to zero.

Patterns of observed eDNA concentrations conform to previous knowledge of *Pycnopodia* ecology. Here we found subtidal sites had higher concentrations of eDNA than intertidal sites. We also found that Calvert Island intertidal sites had higher eDNA concentrations than Quadra Island intertidal sites. And within subtidal sites, fjords had higher concentrations than the outer islands (Figure 8). One-way ANOVA confirms a significant effect of region on eDNA concentration, F(3,236) = 8.323, p <0.0001. Tukey HSD post-hoc results show that all pairwise comparisons with the Subtidal Fjord region are significant with p < 0.0001. The subtidal samples from the fjords near Calvert Island had an average concentration of *Pycnopodia* DNA of 69,941 copies/L (SD = 153544 copies/L) while the subtidal samples from the outer islands sites around Calvert Island had an average DNA concentration of 5943 copies/L (SD = 13,067 copies/L). The intertidal sites on Calvert Island had a similar mean DNA concentration of 3,490 copies/L (SD = 8,403 copies/L). However, this mean is driven by one timepoint from North Beach with a concentration of 30,407 copies/L. With it removed, the Calvert Island intertidal samples then have a mean concentration of 1,433 copies/L. Quadra Island intertidal sites have the lowest concentration of eDNA (mean = 33 copies/L, sd = 112 copies/L, max = 424 copies/L) with the majority of values below the 200 copies/L LoQ for this assay. We also note that when using the raw concentrations not filtered below the LoD, both intertidal surveys had a higher average coefficient of variation (mean = 1.55 ± 0.367, mean ±SD) than the subtidal surveys (0.976 ± 0.522) and a higher frequency of non-detections across bottle replicates within intertidal sites (52.9%) and subtidal sites (1.67%) (See Supplemental Materials).

**Figure 8.**
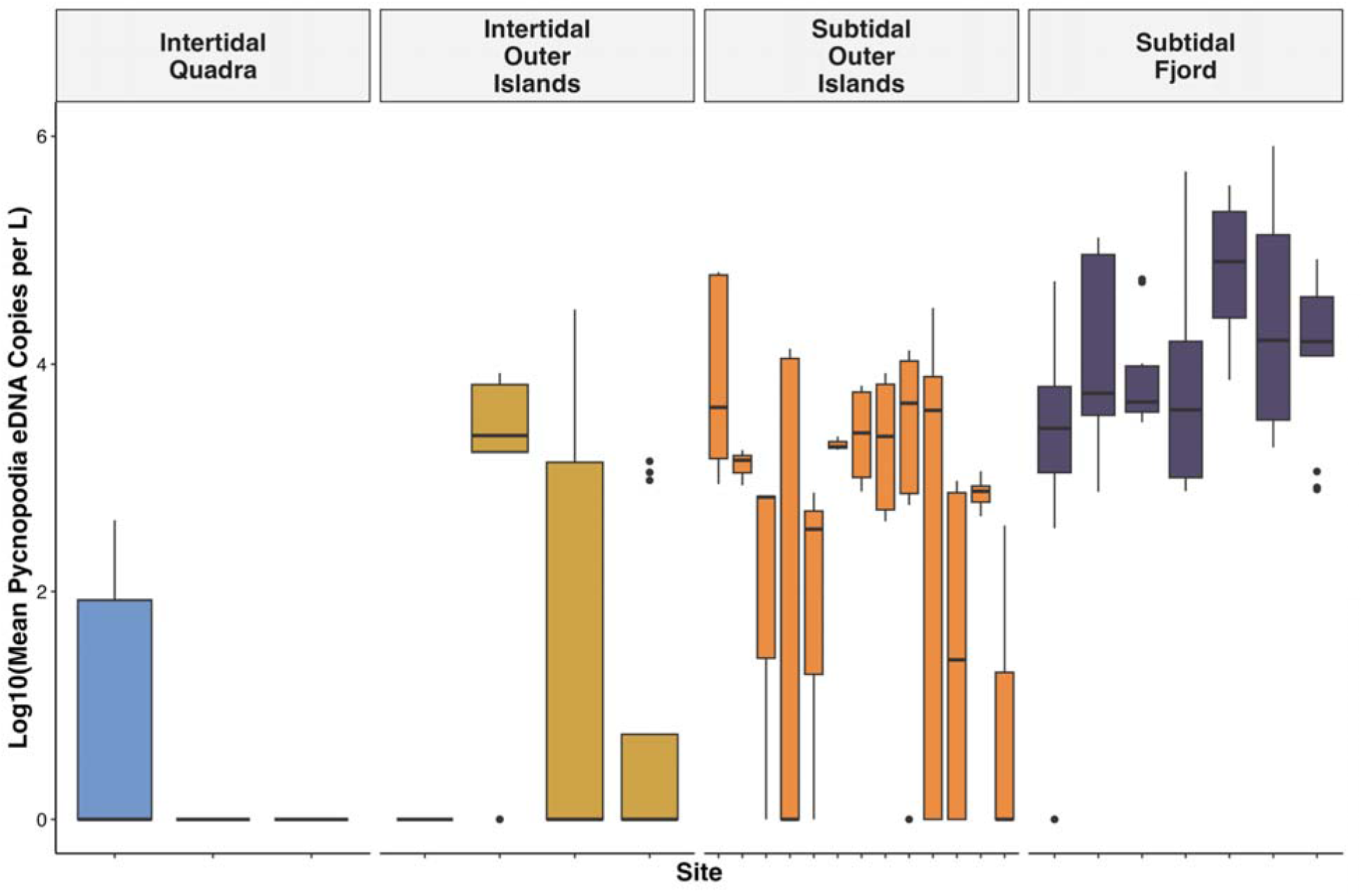
Log10 transformation of copies per liter of *Pycnopodia* eDNA detected in water samples by site. Samples below LoD were set to zero. Colors correspond to the region the site is located within. Each box represents the interquartile range spanning from the 25th to the 75th percentile with the line inside the box marks the median and vertical lines represent 1.5 times the interquartile range. One-way ANOVA confirms a significant effect of site on eDNA concentration, F(26,213) = 2.963, p <0.0001.

When we compared the log10 transformed DNA concentrations from the ddPCR assay from each bottle to the biomass-density calculations from the visual surveys from all subtidal samples, we see a positive yet weak relationship between *P. helianthoides* observed and *P. helianthoides* eDNA concentrations measured (Figure 9, R^2^ = 0.04, p-value = 0.007). However, we see a stronger correlation between ddPCR assay concentrations and visual survey biomass estimates at the site level (R^2^ = 0.45, p-value = 0.001). We also observe strong correlations at the 5 km averaged (R^2^ = 0.58, p-value = 0.01) and region averaged (R^2^ = 0.68, p-value = 0.044), acknowledging that broader spatial scale averages are increasingly subjected to small sample biases that improve correlations (Cramer, 1987). These results highlight the importance of matching spatiotemporal integration scales between visual surveys and eDNA samples as well as underscoring the need for replication on an appropriate scale to account for inherently variable measurements from both sampling methods.

**Figure 9.**
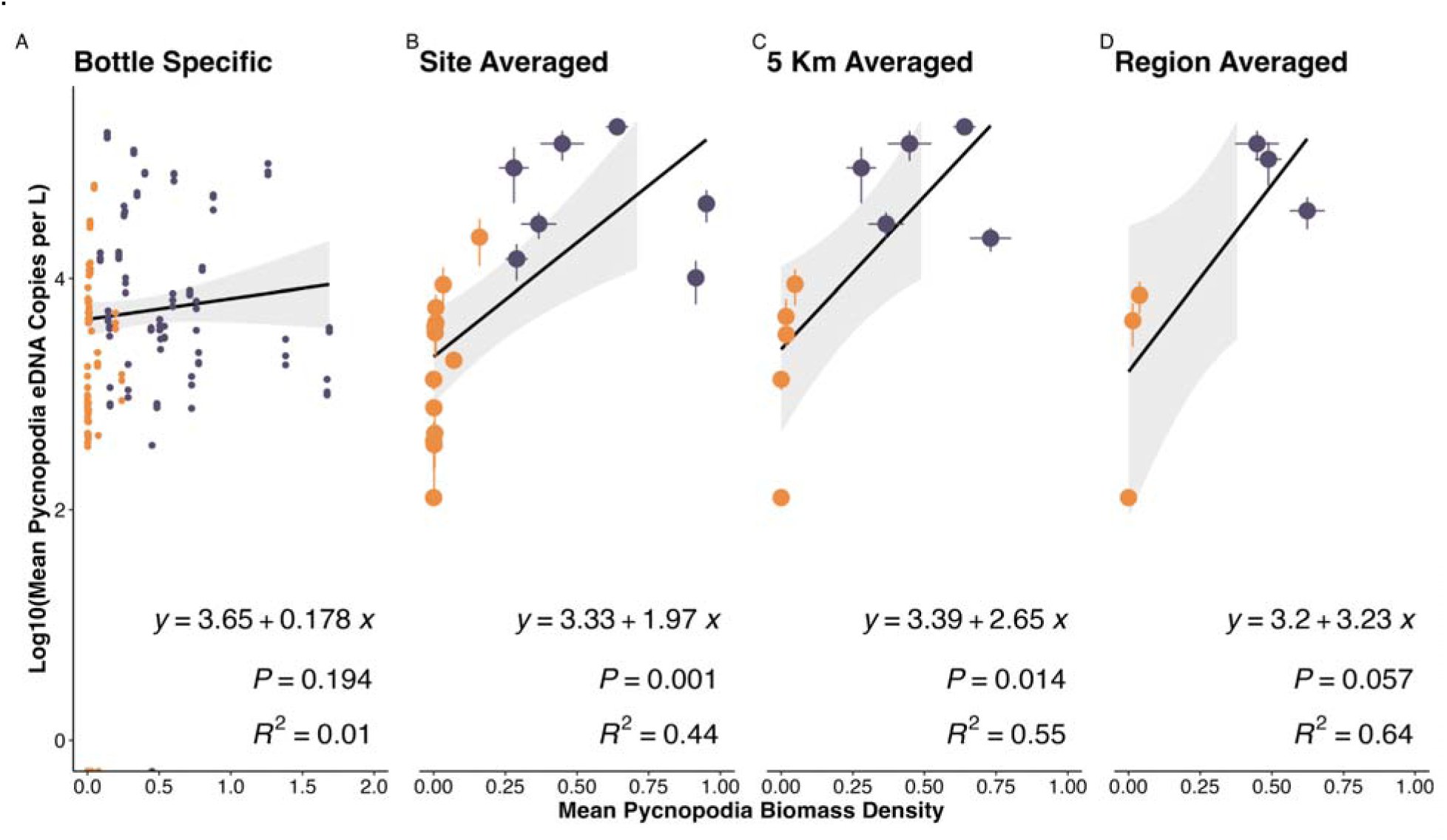
Comparison of eDNA concentration and underwater visual census based *Pycnopodia helianthoides* biomass density estimates across increasing geographic scale. eDNA concentrations measured by ddPCR assay after log10 transformation. Concentrations below the limit of detection were set to zero. *Pycnopodia helianthoides* biomass density at the transect level was calculated as kg per 10 m2 using a length–biomass conversion from Lee et al. 2016. We find better alignment in eDNA and diver abundance estimates when averaging at larger scales as compared to point transect and bottle comparisons. Fjords are depicted in purple and Outer Islands are in orange.

## Discussion

Here we developed and validated a highly specific and sensitive eDNA hydrolysis probe-based PCR assay for the sunflower sea star *Pycnopodia helianthoides*, demonstrating applicability for both monitoring standing populations and identifying rare remnant individuals. We demonstrate a positive relationship between the concentration of *P. helianthoides* DNA copies per L and biomass density in mesocosm and field studies, suggesting that these eDNA assays have potential to estimate abundance at the regional and habitat scale given adequate replication and sampling intensity. Our results highlight the need to better understand the drivers of submesoscale eDNA generation, fate, and transport to accurately interpret eDNA signatures (Andruszkiewicz et al., 2019; Brandão-Dias et al., 2025).

### Assay Sensitivity

The *P. helianthoides* eDNA assay was highly sensitive with a limit of detection and quantification under 1 copy per reaction following the Lesperance et al. 2020 methods for both qPCR and ddPCR approaches (Table 1). We demonstrate the ability to detect as few as a single individual within a 500 m range in coastal marine environments. Previous work has found eDNA approaches to be highly effective at detecting rare species in marine and aquatic environments (Cooper et al., 2022; Hunter et al., 2018, 2026; Langlois et al., 2025; Mauvisseau et al., 2019; Ramón-Laca et al., 2021; Roux et al., 2020). Here we demonstrate similar capabilities for our *P. helianthoides* eDNA assay in nearshore habitats.

A commonly used metric of sensitivity for hydrolysis probe based assays is the limit of blank, limit of detection, and limit of quantification (Bustin et al., 2025). However, there is no consensus within the literature on which approach to calculate LoB, LoD, and LoQ should be applied (Guri, Ray, et al., 2024; Klymus et al., 2020; Lesperance et al., 2021). Here we report both eLowQuant and Klymus et al. 2020 metrics and find a significant difference in sensitivity for both qPCR and ddPCR assays using both methods.

Importantly, we found that the method of LoD and LoQ calculation had a greater role on observed LoD and LoQ (Klymus et al. 2020 versus Lersperance et al. 2021) than between analysis methods (ddPCR or qPCR). Previous work has found that ddPCR approaches are more sensitive than qPCR approaches (Ciesielski et al., 2021; Guri, Ray, et al., 2024). While here we did observe a lower LoD from the ddPCR results from Stanford University (Boehm Lab) than the NOAA PMEL OME qPCR laboratory, the difference was less than 0.1 copies per reaction. In contrast, the calculated LoDs using the Klymus et al. 2021 method were significantly higher, often more than an order of magnitude higher, than those calculated using the Lesperance 2021 approach.

We demonstrate that such discrepancies between LoD and LoQ calculation approaches have real world impact on the interpretation of our field trial results as our observed concentrations from eDNA assays fell between both calculated limits of detection. Clearly these detections are above our LoB (0 copies per reaction) and occurred in multiple technical replicates, strongly suggesting that these are in fact real detections, and thus our eDNA assay is sensitive to detect a handful of translocated and wild individuals (n < 10) within a coastal marine environment. However, these approaches failed to meet the more conservative LoD guidelines set out by Klymus et al. 2020, especially when accounting for LoD and LoQ on a per plate (or batch) basis. Here, such detections of a handful of individuals would require substantially higher replication with greater than 9 technical PCR replicates as well as greater than three 1 L sea water samples. In contrast, our sampling regime was sufficient to detect *P. helianthoides* at both field trials when applying the eLowQuant derived LoD estimates. These results highlight the importance of carefully considering which approaches are used for the LoD and LoQ calculations as these will impact the interpretation of eDNA observations. Ultimately, it is up to the end user of such an assay to determine which statistical approach is appropriate in determining when a sample is above the LoD. Here we highlight that such choices could have implications for determining whether or not low concentrations of *P. helianthoides* eDNA within a sample constitute whether or not the species is present at a site (Figures S5-S14). Such minor statistical choices in the analysis of eDNA data could have substantial management and conservation implications and thus caution is warranted.

Here we also show how the minimum number of accepted droplets as a detection, effectively the determined LoB, has a strong effect on LoD and LoQ. Many researchers have chosen an LoD of three droplets as this can be reliably distinguished from 0 using Poisson distributions (de Kock et al., 2019; Pinheiro et al., 2012; Rare Mutation Detection Best Practices Guidelines, 2020). However, the environmental qPCR and ddPCR fields have come to different determination in calculations of LoD in which most of the qPCR fields rely on standard dilutions curves while ddPCR approaches often rely on droplet enumeration alone. Such LoQ and LoD calculations for qPCR rely on accurate determination of standard concentrations which may not be sufficiently accurate to reliably determine limits under one copy per reaction given the observed variance in vendor supplied synthetic DNA concentrations.

We also note that detection limits and quantification limits will also depend on the volume and dilution of template added to the reactions (Emslie et al., 2019; Košir et al., 2017). Here we demonstrate that increasing the input template DNA in a 20 µL ddPCR reaction volume (e.g. 2 µL at Hakai vs. 5 µL at Stanford) can decrease the LoD and increase the sensitivity of the assay (Table 1). However, it’s important to recognize the challenging balancing act of changing sample input volumes on assay performance and establishing long term eDNA biorepositories. For example, increasing the volume of template may increase the number of target molecules, but it also has the potential for increasing inhibitor concentration. Using a larger volume and increasing the number of replicates also increases the constraints on the total volume of available template DNA, potentially interfering with the desire to conserve some of that for other applications. We also highlight that LoD and LoQ are affected by the number of partitions generated and used for ddPCR approaches as well as the total volume of the reaction (Basu, 2017; Vynck et al., 2023). Researchers should consider all these factors when aiming to maximize sensitivity of a given assay.

We also demonstrate that the number of freeze thaws likely plays a role in our ability to detect rare targets, especially those just above our LoB and at or below the LoD. Here we found an increased number of non-detections from samples that had previously amplified after undergoing three freeze thaw cycles. These results align with findings reported by previous studies (Huge et al., 2022; Mauvisseau et al., 2021). Unfortunately, our highest number of observed amplifications occurred on the first qPCR plate at NOAA PMEL which had poor standard curve performance. By the time we successfully reduced our pipetting error by Pycno Plate Run 5, we had incurred 6 total freeze thaw cycles and could no longer amplify any eDNA from the Friday Harbor Laboratories’ Intake Pipes. These results highlight the need for extreme care when attempting to amplify exceedingly low concentrations of eDNA from few individuals in the field and suggest that robust routines to reduce freeze thaws may need to be implemented to ensure accurate detection of extremely rare targets (e.g. running hydrolysis probe based assays immediately after DNA extraction to avoid any freeze thaw cycles).

We found Sterivex filtration captured an order of magnitude more DNA from sunflower sea star mesocosms compared to passive and Smith Root filter approaches (Supplemental Materials, Figure S4). These results align with previous comparisons of gauze based passive filters and sterivex filters (Kressler et al., 2026). However, we can’t disentangle the effects of preservation and material on eDNA recovery. Future work understanding the interaction of eDNA capture approach and efficacy of preservation strategies is warranted (Andruszkiewicz Allan et al., 2025; Bessey et al., 2021; Chen et al., 2024; Maiello et al., 2024; Spens et al., 2017; Tomke et al., 2025; von Ammon et al., 2023). Regardless, we demonstrate that both gravity filter and peristaltic pump protocols filtering one liter of sea water onto 0.22 µm Sterivex filters are effective at detecting *P. helianthoides* from both mesocosm and field settings.

The eDNA assay sensitivity is exemplified by the consistent detection of *P. helianthoides* from intertidal samples where no *P. helianthoides* individuals were observed. Given that *P. helianthoides* is known to occur at these sites subtidally, these results strongly suggest that surface intertidal samples are capable of detecting subtidal individuals (Gold et al., 2023; Kelly et al., 2018; Shea & Boehm, 2024). Together, our laboratory, mesocosm, and field results strongly suggest that our eDNA assay has the capability to detect even very few *P. helianthoides* in the wild, enabling a wide range of applications for monitoring wild populations and reintroduction efforts.

### Biological Influences on eDNA Signatures

The amount of eDNA present in the environment is in part a function of biological influences including physiology, morphology, behavior, life stage, allometry (Brandão-Dias et al., 2025; Harrison et al., 2019; Shelton et al., 2023; Yates et al., 2021). Here we observed that disease state can also influence apparent DNA generation rates from opportunistic samples from wasting versus non-wasting *P. helianthoides* individuals; these are the first such observations for any species that we are aware of. Given the symptoms of SSWD result in the breaking apart of the tissues, it is not surprising that such individuals release a greater amount of eDNA into the water column (Fuess et al., 2015; Hewson et al., 2014). Further work understanding the influence of SSWD and other diseases on eDNA generation is warranted for accurate interpretation of observed eDNA concentrations.

We found that larval *P. helianthoides* produce a substantial amount of eDNA in mesocosm experiments (Figure 6). Our results suggest that sampling during broadcast spawning events would likely result in the collection of substantial quantities of eDNA, a phenomenon both hypothesized and observed for other marine species (Bylemans et al., 2017; Ip et al., 2023; Thalinger et al., 2019; Tsuji & Shibata, 2025; Vautier et al., 2023). Our work adds to the body of literature highlighting the importance of accounting for variation in life stage being sampled for the accurate interpretation of eDNA results (Crane et al., 2021; Kwong et al., 2021; Ostberg & Chase, 2022; Wilder et al., 2023). Importantly, larval counts would not be included in any diver based visual surveys. Interpretation of eDNA signatures from *P. helianthoides* should consider the effect of broadcast spawning which may increase the likelihood of detecting stars, particularly when in low abundance (Gravem et al., 2021), but also suggests that the accidental capture of a larvae onto a filter could lead to dramatic differences in recovered DNA concentrations.

### Spatiotemporal Integration of eDNA Signatures

Patterns of observed eDNA concentrations aligned well with diver based *Pycnopodia helianthoides* density estimates: both approaches detected higher sunflower sea star abundances within subtidal fjords (Gehman et al., 2025). This pattern was particularly evident at regional habitat scales between 5 and 20 km (Figure 9). In contrast, we observed a weak relationship between observed *P. helianthoides* and measured eDNA concentrations at fine spatiotemporal comparisons of individual one liter sample bottles and biomass density from paired dive transects. These results are expected given the different sampling resolution of each method with divers only surveying 30 m x 2 m per transect while eDNA is integrating signatures across tens to hundreds of meters over both space and time. These results align with previous observations of eDNA transport in the Southern Salish Sea, Johnstone Strait and central coast of BC (Duprey et al., 2023; Kelly et al., 2016, 2018; Millard-Martin et al., 2024; O’Donnell et al., 2017; Robinson et al., 2023; Wallace et al., 2018; Xiong et al., 2025).

Individual adult *P. helianthoides* will act as a point source of eDNA, generating patchy intermittent plumes of eDNA over time that are a function of physiology, life state, allometry, behavior, disease state, and population density. These eDNA pulses will be advectively transported in marine systems as well as influenced by the decay, dilution, and diffusion of eDNA molecules in the environment (Brandão-Dias et al., 2025; Harrison et al., 2019; Li et al., 2026). The rate of decay from each sea star eDNA plume is a function of microbial activity, temperature, and sunlight as well as advective and diffusive transport of eDNA molecules (Andruszkiewicz et al., 2017, 2019; Joseph et al., 2022; McKnight et al., 2024; Salter, 2018). Therefore, the eDNA captured within any given 1 L sea water sample will be a result of underlying species’ biology and demographics as well as fate and transport of eDNA in the environment (Brandão-Dias et al., 2025).

Given these influences on recovered eDNA signatures, when there are few *P. helianthoides* eDNA plumes within a given site, we expect higher variability across samples (Figure S6). This is because many samples will invariably sample outside the handful of rare plumes within a given site, resulting in little to no eDNA detections. Yet we expect a handful of rare samples will be collected directly within a plume, resulting in substantially higher yielded eDNA concentrations. In contrast, in regions with higher sunflower sea star densities, we expect that any given bottle sampled will be within at least one, if not multiple, turbulent plumes of *P. helianthoides* eDNA. We expect this to result in consistently higher recovered eDNA concentrations with lower total variance across sample bottles. Here, we observed higher variance and lower mean concentrations in intertidal sites that had extremely few (n=1) observed intertidal *P. helianthoides* as compared to subtidal sites with substantially higher *P. helianthoides* densities (Figure 8), aligning with the proposed model of both detections and concentrations being a function of underlying sunflower sea star densities and advection and diffusion of eDNA plumes.

In further support of such a framework, we also observed higher variance in both observed DNA concentration and frequency of detection at these intertidal sites with likely unobserved subtidal individuals than from subtidal sites with confirmed *P. helianthoides* occurrences. These results align well with our understanding of the fate and transport of eDNA in marine environments: patchy eDNA signals being less frequently and more variably detected further from the generating source of eDNA (Bessey et al., 2020; Sanches & Schreier, 2020). These results also suggest that the frequency of non-detections across samples is likely a strong indicator of the underlying density of a species in a given habitat sampled; a higher rate of non-detections across sample and PCR replicates being strongly associated with either fewer individuals and/or individuals further away from the sampling location (Supplemental Results). We note that such observations of eDNA suggest strong mechanistic similarities to sensing odor plumes (Koehl et al., 2001; Michaelis et al., 2020). Together, these observations are valuable in the context of interpreting *P. helianthoides* eDNA signatures to identify the location of putative wild individuals, especially as sunflower sea stars remain considerably rare in much of its southern range.

Importantly, our results provide generalizable information on the spatiotemporal integration of eDNA signatures from coastal marine environments. An eDNA sample from a tidally influenced temperate fjord system is likely integrating input DNA across tens of meters to multiple kilometers across a temporal scale of hours to days (Xiong et al., 2025). In contrast, diver based underwater visual census has a much narrower spatiotemporal integration across a 30 m x 2m belt transect completed in under 30 minutes. It is therefore unsurprising that narrowly paired *in situ* observations with dramatically different spatial and temporal integration have poor correlation (Figure 9). Instead, we found more accurate concordance in observations at the 1, 5, and 20 km scales in which the scale of resolution was likely equal or greater to the spatiotemporal integration of eDNA signatures. We highlight that the higher concordance between methods occurred at a scale of aggregation similar to the scale fate and transport of eDNA molecules in the Southern Salish Sea (Kelly et al., 2018; Xiong et al., 2025), noting that the regions sampled here in the British Columbia’s Central Coast and Northern Salish Sea often have much larger advective transport (MacCready et al., 2020; Pawlowicz et al., 2019). This resulted in averages across replicate eDNA samples and transects that are fully inclusive of the maximum expected spatiotemporal integration signatures. Our results strongly suggest that accurate abundance estimation must account for spatiotemporal integration of eDNA signatures to determine the minimum scale at which abundance differences can be reasonably delineated.

These results emphasize that understanding submesoscale dynamics of eDNA fate and transport within a study system are critical for accurate interpretation of eDNA results (Andruszkiewicz et al., 2019; Brandão-Dias et al., 2025; Xiong et al., 2025). Successful application of eDNA efforts will require accurate estimates of spatiotemporal integration signatures and thus requires accurate estimation of species-specific shedding rates, advective transport, dispersion, and degradation rates (Harrison et al., 2019).

Importantly, this framework then suggests that applying the same sampling schema across disparate marine habitats with different transport regimes may not be appropriate. We highlight here that on the central coast of British Columbia and in the Salish Sea, advective tidally driven transport is greater than many other habitats utilized by *P. helianthoides* along U.S. West Coast (MacCready et al., 2020). We expect that in other coastal systems inhabited by *P. helianthoides* with reduced advective transport, eDNA signatures will differ on the scale of tens to hundreds of meters (Ely et al., 2021; Gold et al., 2021; Monuki et al., 2021; Shea & Boehm, 2024). Such finer scale resolution of eDNA signatures within a low transport environment can enable more precise localization and abundance mapping, but also will require a greater number of total samples within a given area to adequately capture the heterogeneity of eDNA signatures. We highlight that variation in eDNA integration signatures stands in stark contrast to standardized transect based underwater visual census approaches. Taking such information under consideration will be critical for accurate interpretation of a coast-wide eDNA survey to identify existing refugia and monitor the extent of existing *P. helianthoides* populations. Especially as other abiotic and biotic factors beyond advective transport may change across biogeographic regions (e.g. eDNA decay rates given the strong influence of temperature (Lamb et al., 2022)).

For example, samples collected in the Northern Salish Sea will likely have dramatically different spatiotemporal integration scales as compared to samples taken within highly protected sites with minimal wave and tidal wave action (e.g. back side of Channel Islands or East facing reefs along the Monterey Peninsula). Thus, our results add to a growing chorus on the importance of understanding the local factors that influence the spatiotemporal integration of eDNA signatures, particularly eDNA residence time and physical advection transport dynamics, for accurate interpretation of eDNA results in marine and aquatic environments (Kelly et al., 2014).

Estimating spatiotemporal scales for accurate interpretation of eDNA results will require further fate transport studies and models of eDNA in nearshore systems. One promising approach is to leverage controlled reintroduction experiments in which sunflower sea stars are reintroduced in cages within regions with no known individuals to better inform accurate interpretation of eDNA signatures in coastal marine ecosystems (Baetscher et al., 2024; Murakami et al., 2019). Establishing baselines of no known individuals is challenging, but can be based on repeated surveys from existing long term ecological monitoring programs in well characterized regions (Freiwald et al., 2024; Menge et al., 2019). In addition, shedding and decay rate studies are needed to understand the baseline generating and degradation rates in marine ecosystems as well as the biotic and abiotic factors that influence such rates to provide bounding information on parameters that influence the spatiotemporal scale of eDNA signatures (Sassoubre et al., 2016; Waters et al., 2024). Other low cost solutions may involve the use of drifters to inform submesoscale dynamics within nearshore coastal regions (Gonçalves et al., 2019; Ohlmann et al., 2017). Furthermore, incorporating eDNA particle distribution into submesoscale transport models across sampling regimes will also improve the interpretation of eDNA signatures (Brasseale et al., 2025; Xiong et al., 2025). Future work to inform our understanding of the controls and scale of spatiotemporal integration scales of eDNA signatures are critical for implementing effective eDNA survey designs as well as accurate interpretation of eDNA signatures.

Despite the need for accurate delineation of spatiotemporal eDNA signatures, our results suggest that with sufficient replication across space and time as well as appropriate concordance of spatiotemporal survey scale, eDNA approaches can capture meaningful differences in *P. helianthoides* prevalence and biomass at regional scales. These results align with previous research that demonstrate eDNA approaches can provide meaningful abundance estimates (Baetscher et al., 2025; Guri et al., 2025; Guri, Shelton, et al., 2024; Shelton et al., 2022; Spear et al., 2021), particularly when well calibrated to local transport conditions and species specific contexts (Parsley et al., 2025; Thalinger et al., 2021).

### Value of Reference Mitogenomes for eDNA qPCR Assay Design

The generation of a species specific qPCR assay was greatly enhanced by efforts to generate the most comprehensive Asteriid mitochondrial genome database to date (Jossart et al., 2024; Sun et al., 2022). By doing our due diligence to capture both intraspecific variation across our target species range and interspecific variation across known congeners, we were able to utilize existing bioinformatics tools to identify conserved *Pycnopodia helianthoides* genes that serve as the template for qPCR assay design (Lopez et al., 2025). Often qPCR assay design is fraught with difficulties in large part because of poor coverage of both target and congener sequences (Langlois et al., 2021). Here we demonstrate that putting in significant *a priori* effort to develop comprehensive mitogenome databases dramatically eases assay development and enhances success (Goldberg et al., 2016; Klymus et al., 2020).

Furthermore, we echo the calls of researchers highlighting for the need for generating full mitochondrial genomes for future barcoding efforts to enhance eDNA assay design (Hoban et al., 2022; Quattrini et al., 2024; Trevisan et al., 2019). Here our assay targets an *nad5* gene that is an extremely rare barcoding gene with a paucity of available sequences on public reference databases. However, the *nad5* gene is almost entirely unique and conserved within *P. helianthoides* making an excellent target for assay development. Mitogenomes produce upwards of twenty times the data as a typical single gene barcode at only a few times more the cost of capillary sequencing (Gold et al., 2025). However, we argue that such differences in price are well worth the time saved in the assay development and design and provide more robust genetic resources that enable a broader range of science beyond species identification (Allison et al., 2023; Langlois et al., 2025). Further work and efforts to develop comprehensive and curated regional reference barcode databases such as the California Intertidal DNA Barcoding Library Project are needed to enhance our ability to efficiently and effectively develop eDNA assays for species of interest (Crowfoot et al., 2024).

### Towards FAIR qPCR eDNA Data Mobilization

Detailed and accurate reporting of sample collection, nucleic acid extraction, assay design, cycling conditions, and bioinformatic analysis of qPCR protocols is required for reproducibility, accurate quantification, technological progress, and reliable diagnostic or research applications (Borchardt et al., 2021; Bustin et al., 2025). Yet, targeted eDNA methods are often not reported in a manner that meets these needs; protocols are not always findable, accessible, interoperable, and reusable (FAIR) (Wilkinson et al., 2016). To address this, we provide detailed standard operating procedures for running qPCR and ddPCR single species assays adhering to FAIRe standards (Takahashi et al., 2025) and Better Biomolecular Ocean Practices (BeBOP) guidelines (Samuel et al., 2021). This work builds on (Pitz et al., 2026) (submitted) to enable the implementation of the Environmental Microbiology Minimum Information (EMMI) and Minimum Information for Publication of Quantitative Real-Time PCR Experiments (MIQE) guidelines in a standardized, machine-readable Markdown format to enhance FAIR data delivery (Borchardt et al., 2021; Bustin et al., 2025). This work enhances FAIRe and BeBOP efforts, both endorsed Projects under the UN Decade of Ocean Science for Sustainable Development Ocean Biomolecular Observing Network (OBON) (Leinen et al., 2022), with resources for *P. helianthoides* that can be easily adapted for other targeted eDNA assay deployment. These efforts more easily allow protocol details—from extraction to data analysis—to be shared using standardized and interoperable terminology, making these data machine-readable and artificial intelligence (AI)-ready. *P. helianthoides* FAIRe BeBOP protocols are provided (https://doi.org/10.5281/zenodo.19711947).

### Application of eDNA Assay for Conservation and Management

Here we present the validation and benchmarking of eDNA assay for *P. helianthoides* sunflower sea stars demonstrating an operational level 5 on the Thalinger et al., 2021 scale equivalent to a NASA technology readiness level 9 (Olechowski et al., 2020) for monitoring sunflower sea stars from coastal environments. We highlight a suite of future applications of the eDNA assay for conservation and management of this iconic echinoderm.

This assay can assist with the evaluation of population recovery status trends and identify the extent and type of critical habitat being utilized by *P. helianthoides* (NOAA, 2023). In particular, coast-wide applications of eDNA efforts can identify remnant intact populations, particularly in Southern Oregon and California where a handful of individuals have recently been spotted for the first time in 2025 in over a decade since the onset of SSWD and the associated mass mortality event (Harvell et al., 2019). Furthermore, as we demonstrate here, eDNA approaches provide a simple and cost effective tool to monitor reintroduced populations. This assay can also be used to interrogate archived samples of eDNA such as the CalCOFI zooplankton net tow biorepository to better understand changes in spawning stock biomass and recruitment dynamics before, during, and after the onset and aftermath of the SSWD pandemic. Preliminary metabarcoding work from the CalCOFI archives has found declines in *P. helianthoides* and other Asteriid star recruitment in the aftermath of the 2014-16 marine heatwave (Gold unpublished). The sensitivity of this assay to detect as few as a single *P. helianthoides* individual within a 500m swath of marine habitat, provides a powerful tool for being able to map and monitor this ecologically important species of conservation concern.

