## Supplemental Materials for "Molecular Star Gazing: Development and Validation of an Environmental DNA Assay for the Imperiled Sunflower Sea Star (*Pycnopodia helianthoides*)"

All supplemental figures are available in high resolution stand alone files at <https://github.com/marinednadude/Pycnopodia_eDNA_assay_validation_MS>

### **Assay Specificity**

#### Results

*Contamination of Patiria miniata tissue*

We observed amplification in one tissue of *Patiria miniata* which was unexpected given the high number of mismatches in the forward (n=10), reverse (n=5), and probe (n=10). No other tissues which had fewer mismatches amplified, including *P. helianthoides* closest relative *Rathbunaster californicus.* In response to the amplification we sourced a new specimen of *Patiria miniata* and observed no amplification. Our assay is highly sensitive to only a few copies of target DNA. Tissue samples that were obtained by NOAA PMEL OME followed proper sampling procedures, but were processed in laboratories and museums that also contain specimens of *P. helianthoides.* Highly conservative sterile techniques which aim to ensure zero contamination of DNA, like those employed in ancient DNA laboratories and other eDNA facilities, are not common practices in routine specimen voucher campaigns. It is also possible that when handling tissue extractions in the laboratory an unintentional mistake was made for this specific sample that was never repeated across no template controls in this study. The most parsimonious explanation in light of this context of high rates of mismatches and lack of amplification in any other tissues or negative controls is that this was a sample specific contamination event.

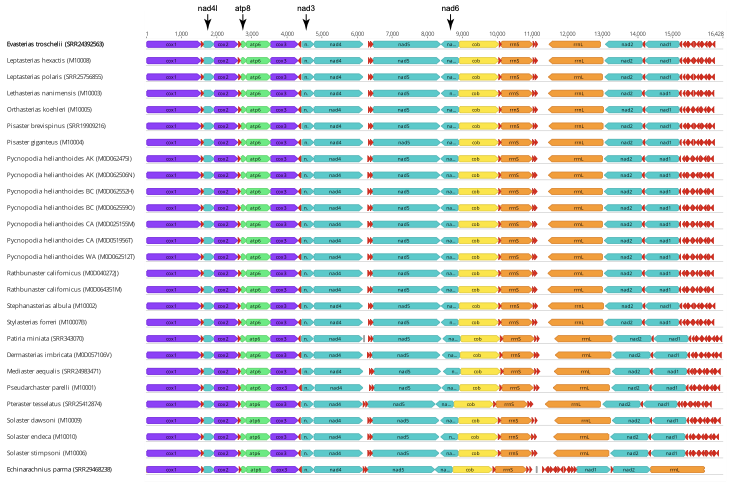

**Figure S1. Alignment of novel mitochondrial genomes generated for this study.**

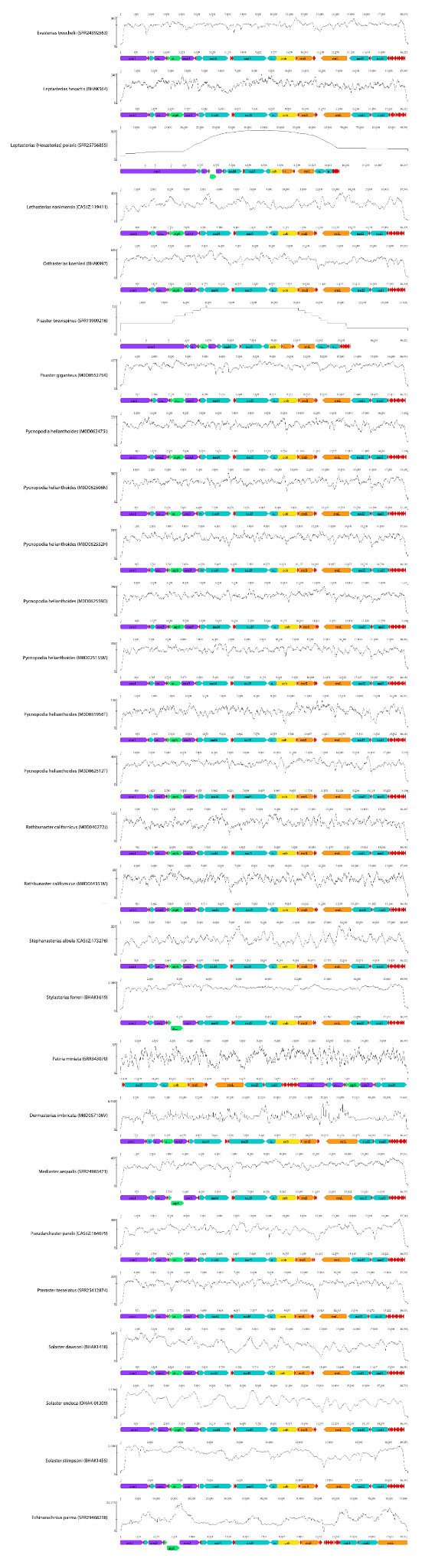

**Figure S2. Coverage plots of novel mitochondrial genomes generated for this study.**

**Figure S3. Alignment of *Pycnopodia helianthoides* nad5 assay.**
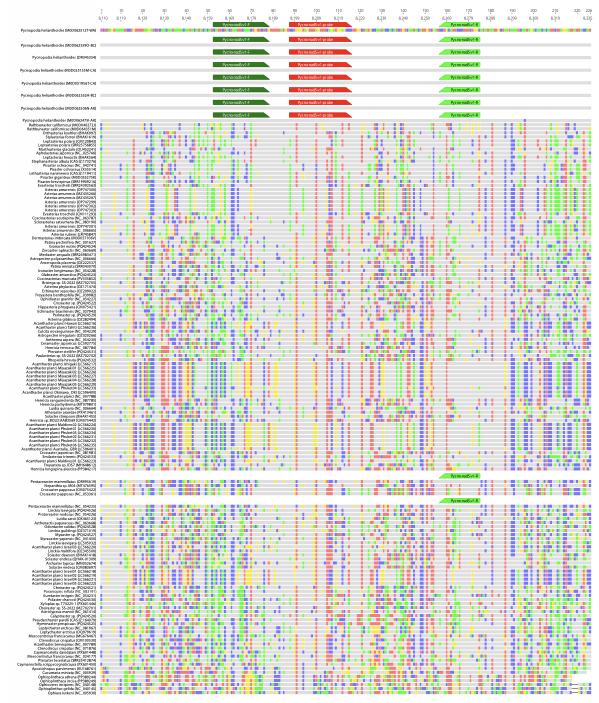

We aligned the designed *P. helianthoides* hydrolysis probe assay against 145 sea stars in the Family Asteroidea and echinoderm outgroups. See Table S12 for summarized counts.

### **Active Versus Passive Filtration**

#### Methods

We collected multiple eDNA samples using different collection approaches: (1) peristaltic pressure filtration of one liter using a Smith Root PES 47 mm diameter 0.22 µm pore size filter (<https://doi.org/10.5281/zenodo.17655098>), (2) peristaltic pressure filtration of one liter through a Sterivex PES 0.22 µm in line filter (<https://doi.org/10.5281/zenodo.17655027>), and (3) passive sampling of eDNA using Kringle Gauze (sterile), a Paint Roller (non-sterile), Vivecare Gauze (sterile), and a Tische environmental filter (non-sterile) placed in a sterile plastic tub in the effluent for 1 hour. Sterivex were preserved with 95% molecular grade ethanol and then immediately frozen at -20C before extraction 6 days later (Table SXX). Passive filters were immediately frozen at -20C. Smith Root filters were self preserved at room temperature following Thomas et al. 2020. Passive filters and Smith Root filters were extracted 7 months later (Table SXX).

Nucleic-acids were extracted from all filters obtained from mesocosm and field eDNA samples at NOAA PMEL using a modified Qiagen Blood and Tissue Kit (sterivex extraction protocol:<https://zenodo.org/records/17655148>; Smith Root extraction protocol: <https://zenodo.org/records/17655213>). The passive filter extraction protocols followed the Smith Root protocol with the addition of a 1 minute Fastprep bead beating step using Zymo ZR BashingBead Lysis Tubes (0.1 & 0.5mm). Negative controls for passive filters consisted of briefly exposing materials to air for 30 seconds before being placed in sterile ziplock bags and then processed the same as the other passive filters. We note sterivex filters were extracted on May 22nd, 2025 while all other samples were extracted on December 12th, 2025.

#### Results

We observed ~2 million copies per L of Pycnopodia target DNA in the Friday Harbor Aquarium discharge water. The discharge is from a flow through system with 39 individual stars of at least 40 cm in diameter. Sterivex 0.22 µm filters yield the highest concentration of Pycnopodia DNA (Figure 5).

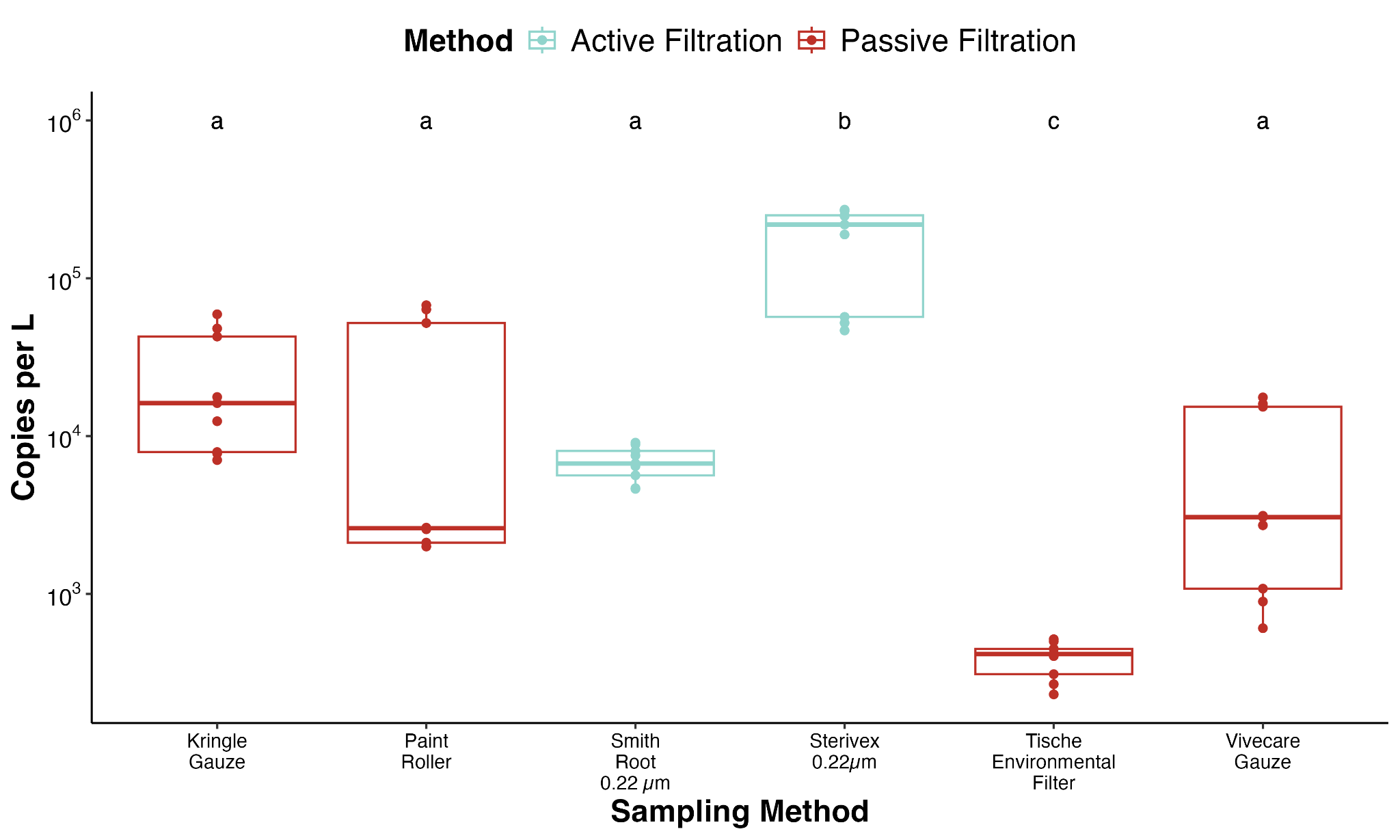

###### **Figure S4. Comparison of sunflower star eDNA yield from mesocosm experiments.** Sterivex 0.22 µm pore size filters yielded significantly more sunflower star from the aquarium discharge tank than Smith Root or passive filters.

#### Discussion

We found Sterivex filtration captured an order of magnitude more DNA from sunflower star mesocosms compared to passive and Smith Root filter approaches. This was surprising given that Sterivex has a smaller surface area (10cm^2) than the Smith Root 47mm filter (17.3 cm^2). These results suggest that differences in the housing or sample integrity from desiccation based preservation are driving these apparent differences. We note that sterivex filters were immediately preserved with 100% molecular grade ethanol in the field before being placed on ice and frozen within 30 minutes of sample collection. In contrast, Smith Root were self preserved via desiccation and left at room temperature for 6 months prior to extraction following the manufacturer’s protocols (Thomas et al., 2019).

Importantly, these results highlight that passive filters can be used for field applications to detect sunflower stars, albeit at a significant disadvantage with even the best performing materials capturing less than a 10th of the DNA captured by active sterivex filtration methods. These results align with previous comparisons of gauze based passive filters and sterivex filters (Kressler et al., 2026). However, we note that we can’t rule out with our study design that preservation of filters in the field is driving these results as passive filters were frozen immediately in the field at -20˚C, but not extracted for 6 months. Future work understanding the efficacy of preservation strategies is warranted (Andruszkiewicz Allan et al., 2025; Spens et al., 2017; Tomke et al., 2025; von Ammon et al., 2023).

Despite these observations, we highlight that passive eDNA approaches can recover *P. helianthodies* eDNA signatures. Given the ease of deployment, passive filtering approaches provide a scalable approach to expand sample coverage and enable citizen and community science participation at a broader scale (Bessey et al., 2021; Chen et al., 2024; Maiello et al., 2024). However, our results highlight that such efforts are better suited for monitoring existing populations with larger standing stock biomass or monitoring reintroduction efforts than for trying to detect rare individuals at the edge of their known range given the order of magnitude difference in observed eDNA recovery efficiency.

Here we highlight that our gravity filter and peristaltic pump protocols filtering one liter of sea water onto 0.22 µm sterivex filters are effective at detecting *P. helianthoides* from both mesocosm and field settings.

### **Paired with visual surveys with No Filtering Applied**

#### Results

Here we report the results from the raw ddPCR in which any amplification was considered detection. No filtering was performed using the limit of detection.

##### Pycnopodia eDNA Detections

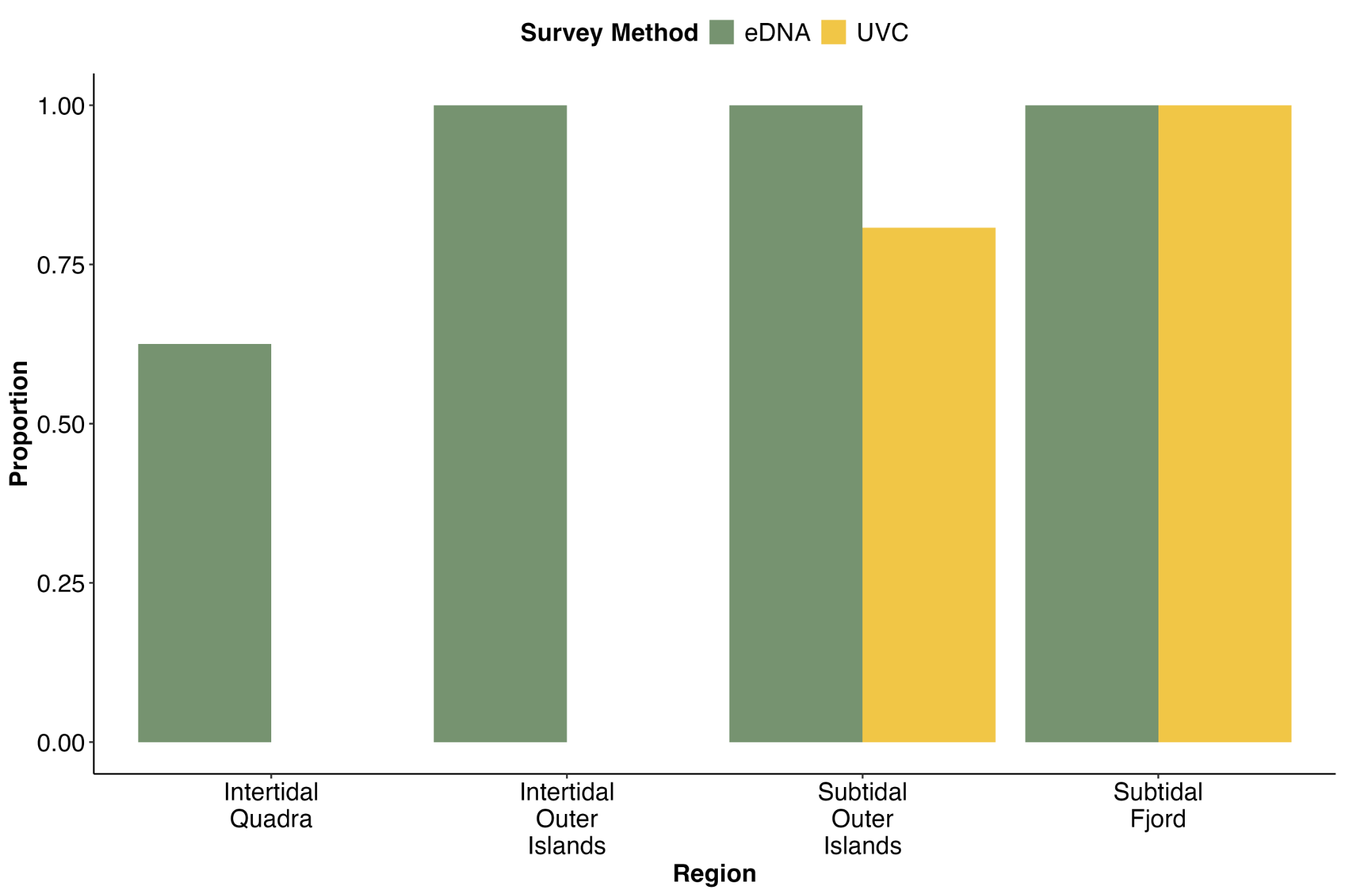
**Figure S5. Comparison of eDNA and diver *P. helianthoides* detections.** Proportion of samples collected from the four regions with a positive detection of *Pycnopodia* by either underwater visual census (UVC) or by ddPCR assay of water samples (eDNA). eDNA detected sunflower sea stars at all subtidal sites as well as a greater proportion of intertidal sites when not applying an LoD filter.

##### Pycnopodia eDNA Concentration Variability

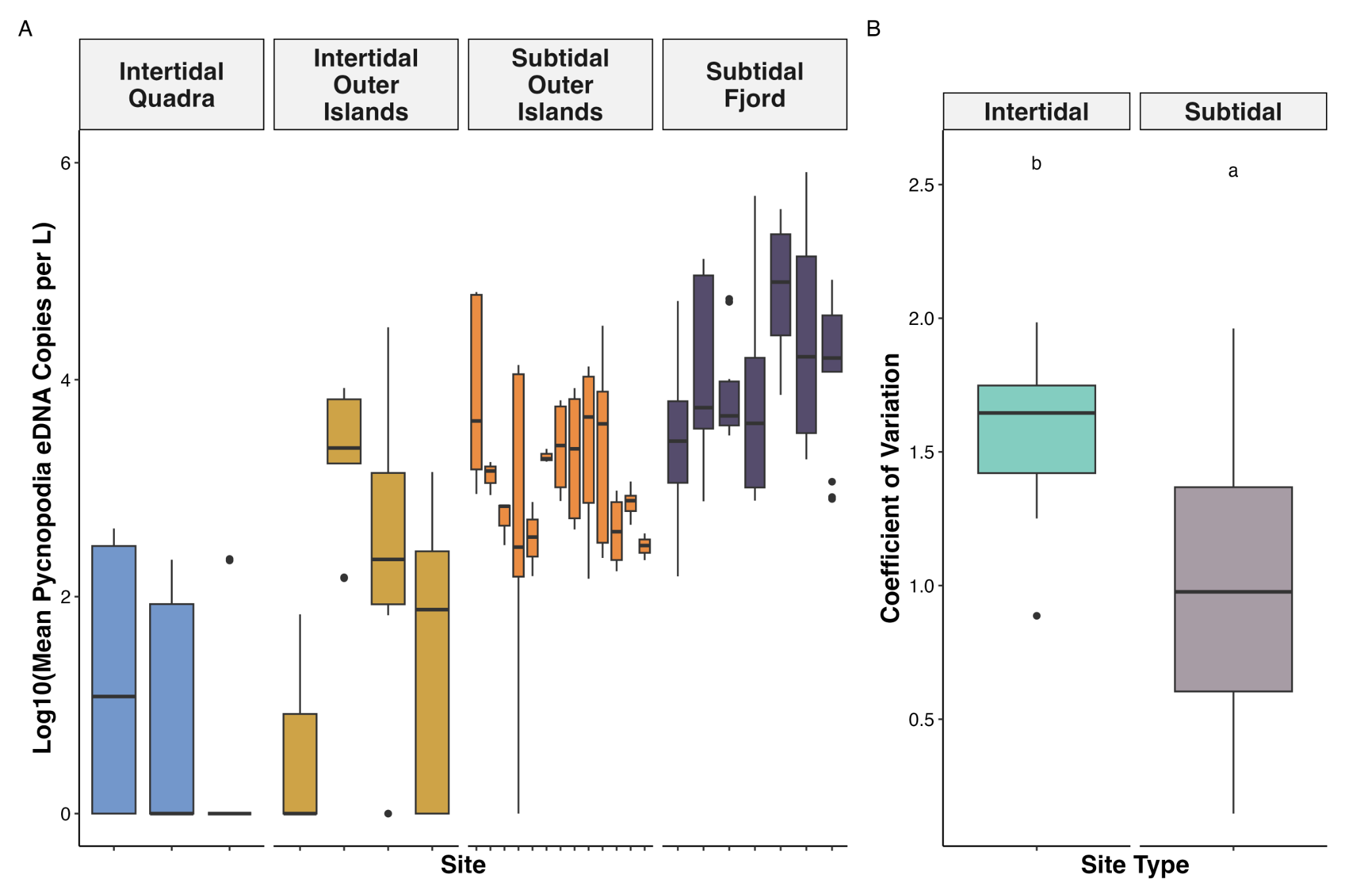

**Figure S6**.  **Variability in captured eDNA concentration across sites.** Log10 transformation of copies per liter of *Pycnopodia* eDNA detected in water samples by site (A). Colors correspond to the region the site is located within. Each box represents the interquartle range spanning from the 25th to the 75th percentile with the line inside the box marks the median and vertical lines represent 1.5 times the interquartile range. We only observed the higher coefficient of variation in intertidal sites when not applying the LoD filtering as many of the low concentration replicates were filtered out. Our results suggest that a high coefficient of variation in raw eDNA observations are likely indicative of low biomass within a site.

##### eDNA Concentration Versus Biomass Density

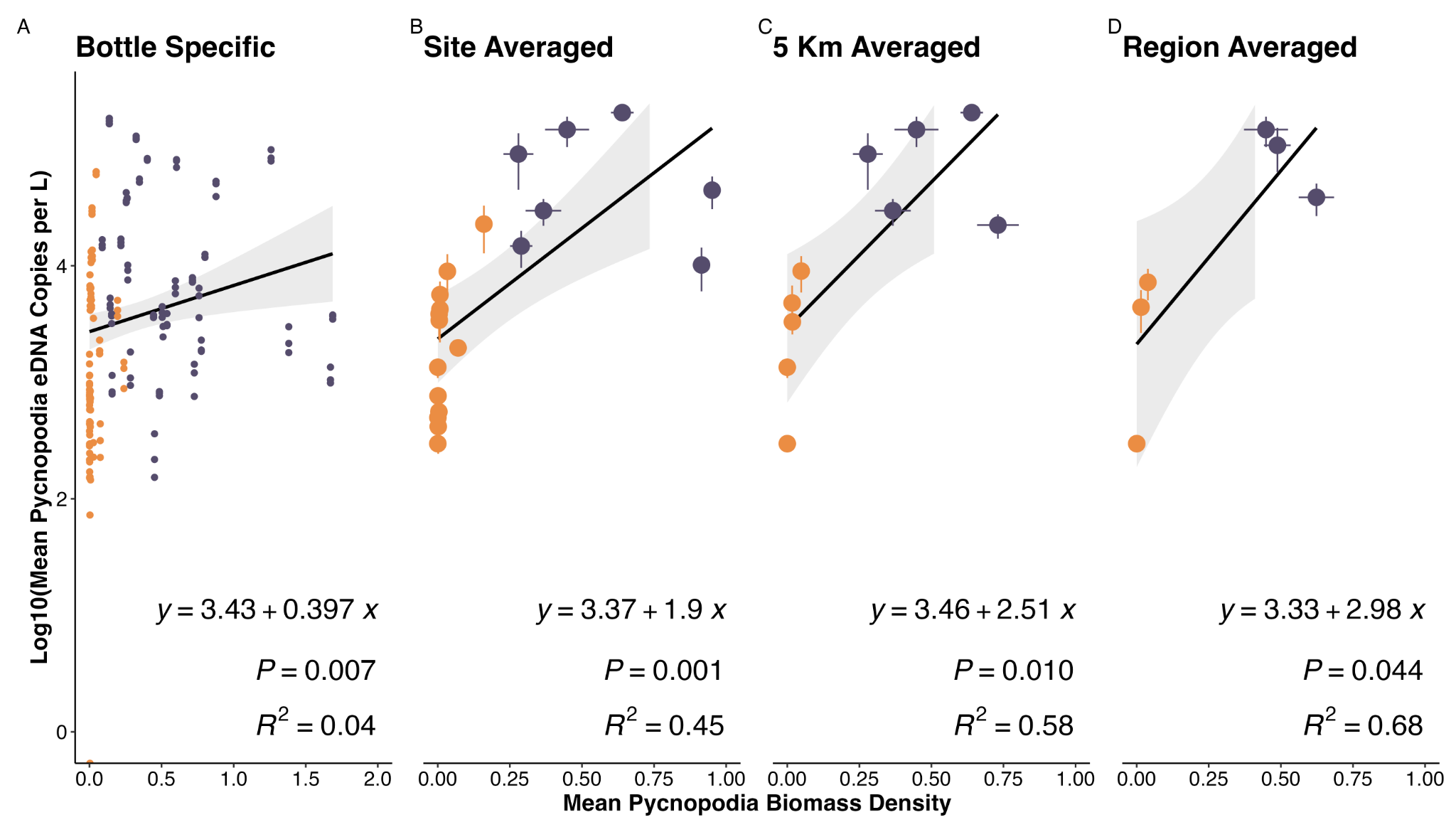

.

**Figure S7**. **Comparison of eDNA concentration and underwater visual census based pycnopodia biomass density estimates across increasing geographic scale.** eDNA concentrations measured by ddPCR assay after log10 transformation. *Pycnopodia* *helianthoides* biomass density at the transect level was calculated as kg per 10 m2 using a length–biomass conversion from Lee et al. 2016. We find better alignment in eDNA and diver abundance estimates when averaging at larger scales as compared to point transect and bottle comparisons.

##### eDNA Concentration Versus Density

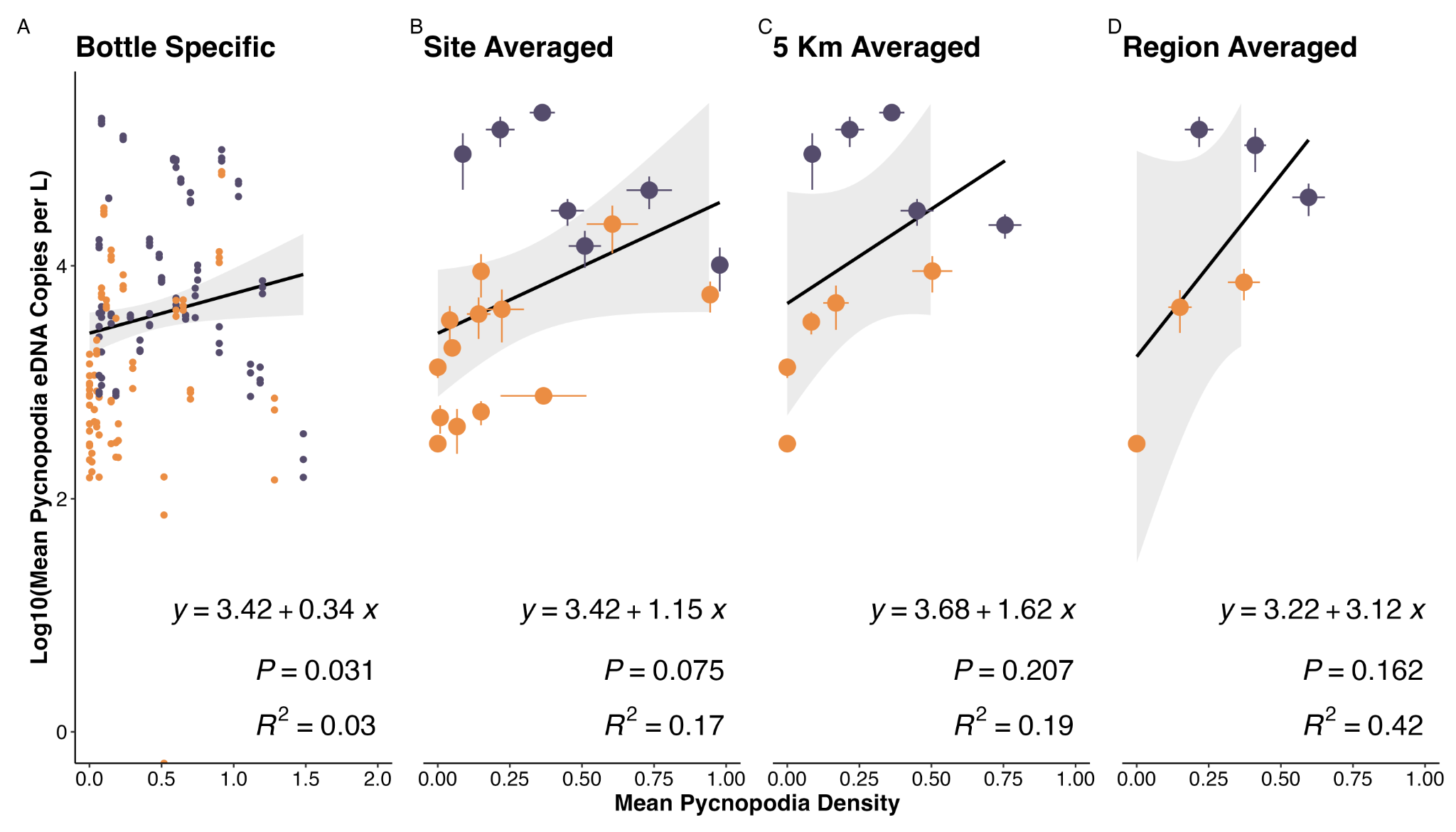

**Figure S8**. **Comparison of eDNA concentration and underwater visual census based pycnopodia density across increasing geographic scale.** eDNA concentrations measured by ddPCR assay after log10 transformation. *Pycnopodia* *helianthoides* density at the transect level was calculated as individuals per 10 m2. We find better alignment in eDNA and diver abundance estimates when averaging at larger scales as compared to point transect and bottle comparisons. We also find better alignment with eDNA biomass density estimates, which account for individual size (Figure 1) than with density alone.

##### eDNA Concentration Versus Counts

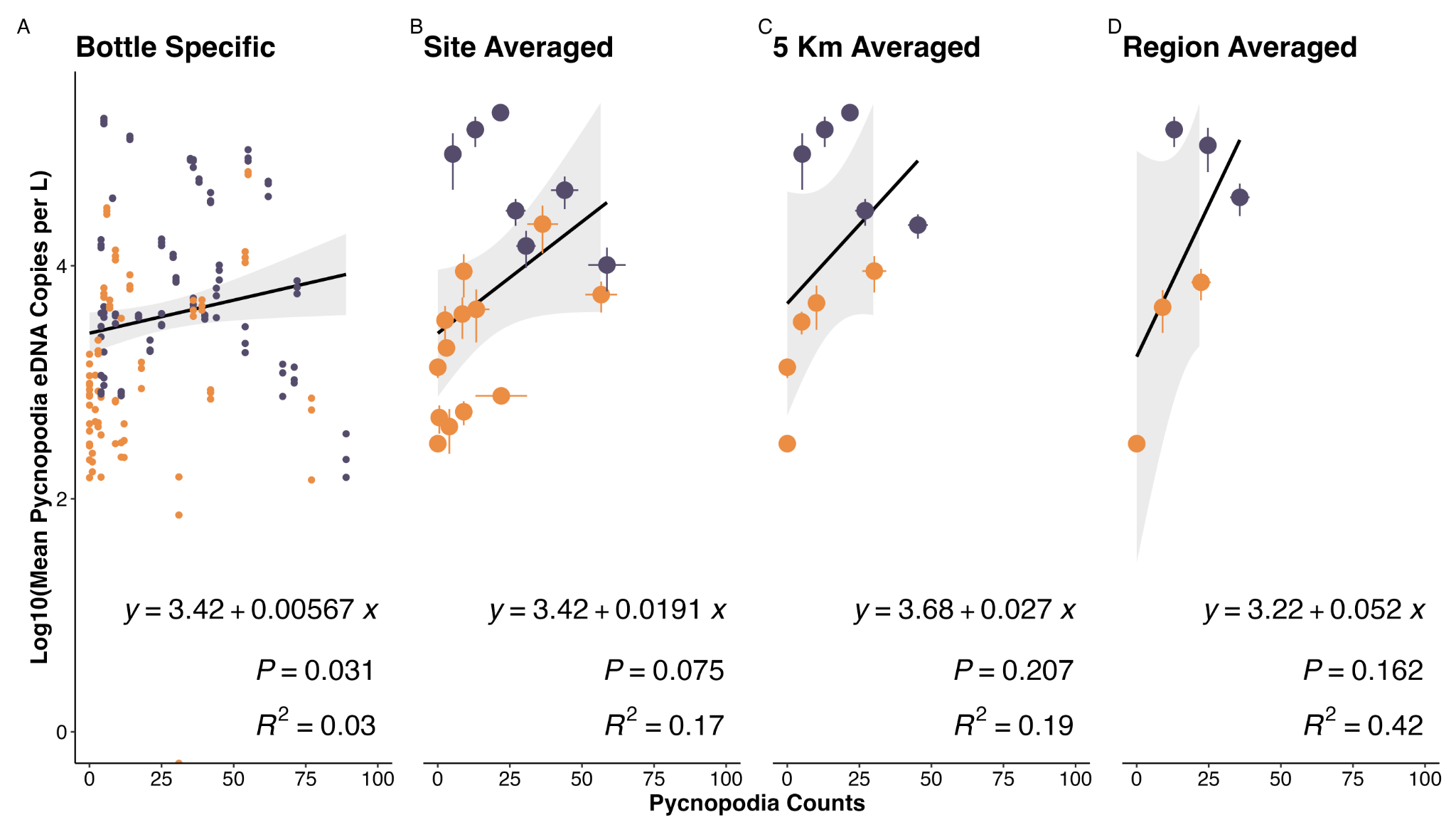

**Figure S9**. **Comparison of eDNA concentration and underwater visual census based pycnopodia counts across increasing geographic scale.** eDNA concentrations measured by ddPCR assay after log10 transformation. We find better alignment in eDNA and diver abundance estimates when averaging at larger scales as compared to point transect and bottle comparisons. We also find better alignment with eDNA biomass density estimates, which account for individual size (Figure 1) than with counts.

### **Paired with visual surveys with Lod Filtering Applied**

#### Results

These results are reported in the discussion section of the manuscript.

##### Pycnopodia eDNA Detections with LoD Filtering

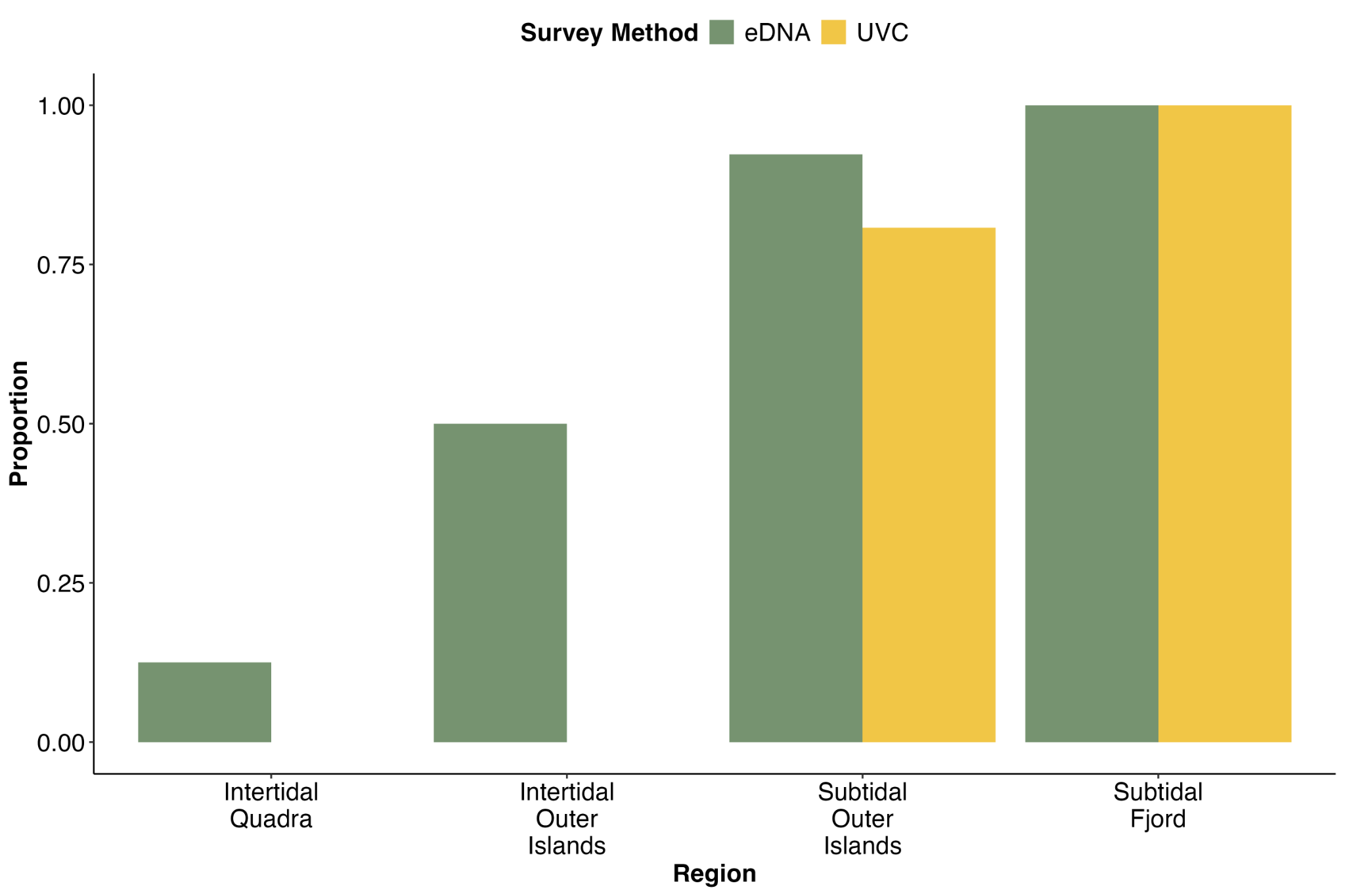

###### **Figure S10. Comparison of eDNA and diver *P. helianthoides* detections with LoD filtering.** Proportion of samples collected from the four regions with a positive detection of *Pycnopodia* by either underwater visual census (UVC) or by ddPCR assay of water samples (eDNA). Samples below LoD were set to zero.

##### Pycnopodia eDNA Concentration Variability with LoD Filtering

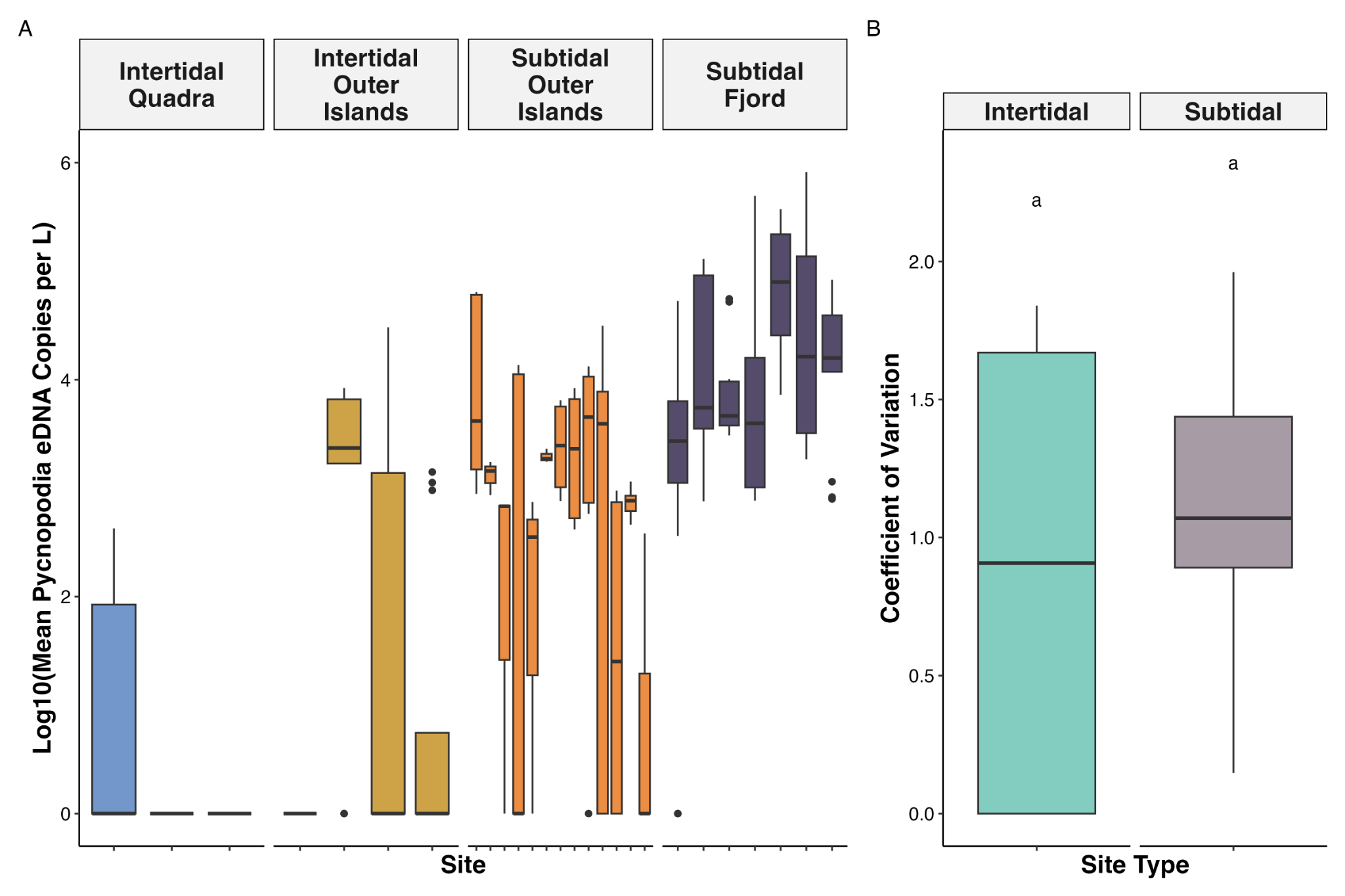

**Figure S11**. **Variability in captured eDNA concentration across sites with LoD filtering.** Log10 transformation of copies per liter of *Pycnopodia* eDNA detected in water samples by site (A). Samples below LoD were set to zero. Colors correspond to the region the site is located within. Each box represents the interquartile range spanning from the 25th to the 75th percentile with the line inside the box marks the median and vertical lines represent 1.5 times the interquartile range.

##### eDNA Concentration Versus Biomass Density with LoD Filtering

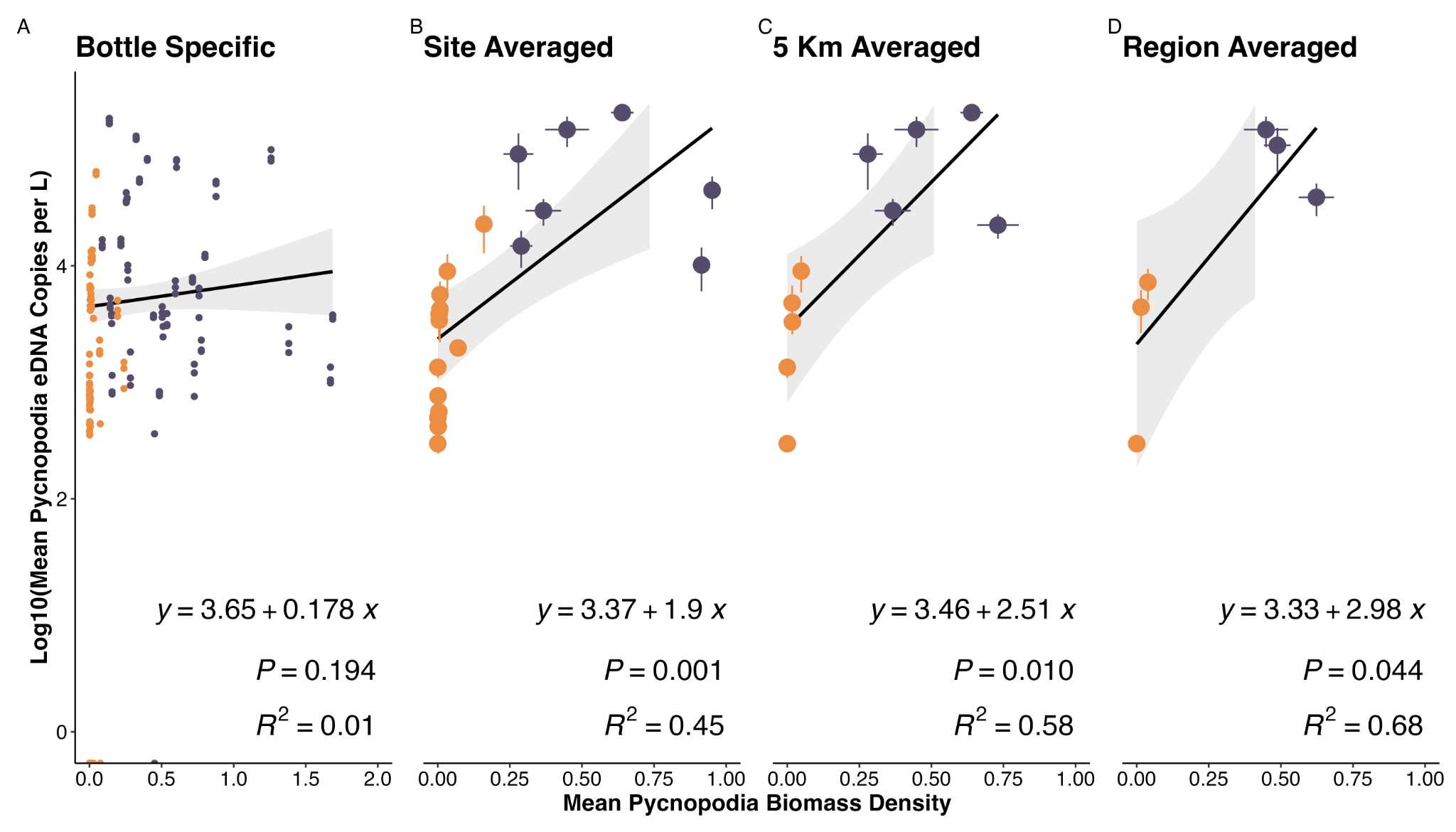

**Figure S12**. **Comparison of eDNA concentration with LoD filtering and underwater visual census based pycnopodia biomass density estimates across increasing geographic scale.** Samples below LoD were set to zero. eDNA concentrations measured by ddPCR assay after log10 transformation. *Pycnopodia* *helianthoides* biomass density at the transect level was calculated as kg per 10 m2 using a length–biomass conversion from Lee et al. 2016. We find better alignment in eDNA and diver abundance estimates when averaging at larger scales as compared to point transect and bottle comparisons.

##### eDNA Concentration Versus Density

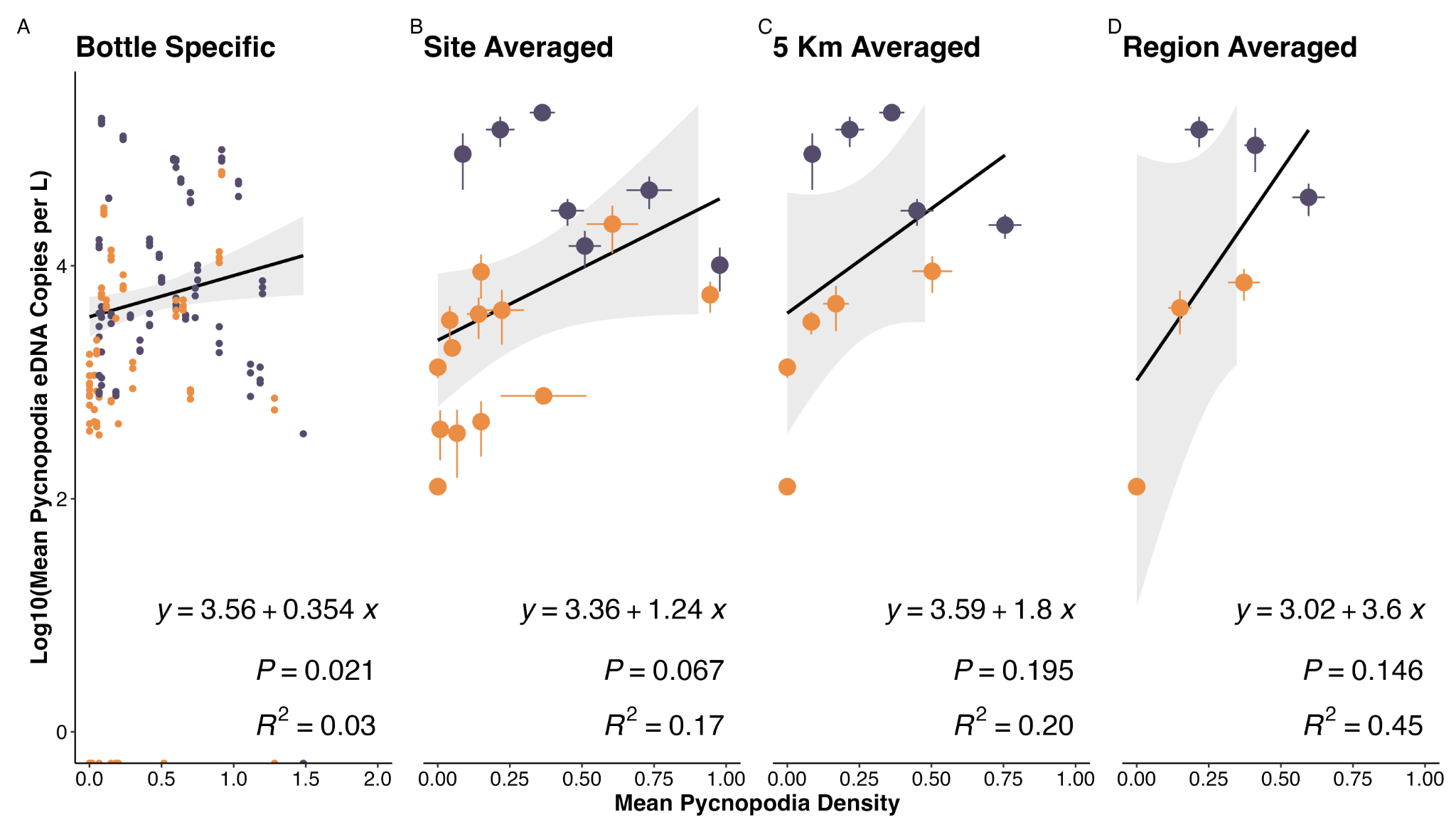

**Figure S13**. **Comparison of eDNA concentration with LoD filtering and underwater visual census based pycnopodia density across increasing geographic scale.** Samples below LoD were set to zero. eDNA concentrations measured by ddPCR assay after log10 transformation. *Pycnopodia* *helianthoides* density at the transect level was calculated as individuals per 10 m2. We find better alignment in eDNA and diver abundance estimates when averaging at larger scales as compared to point transect and bottle comparisons. We also find better alignment with eDNA biomass density estimates, which account for individual size (Figure 1) than with density alone.

##### eDNA Concentration Versus Counts

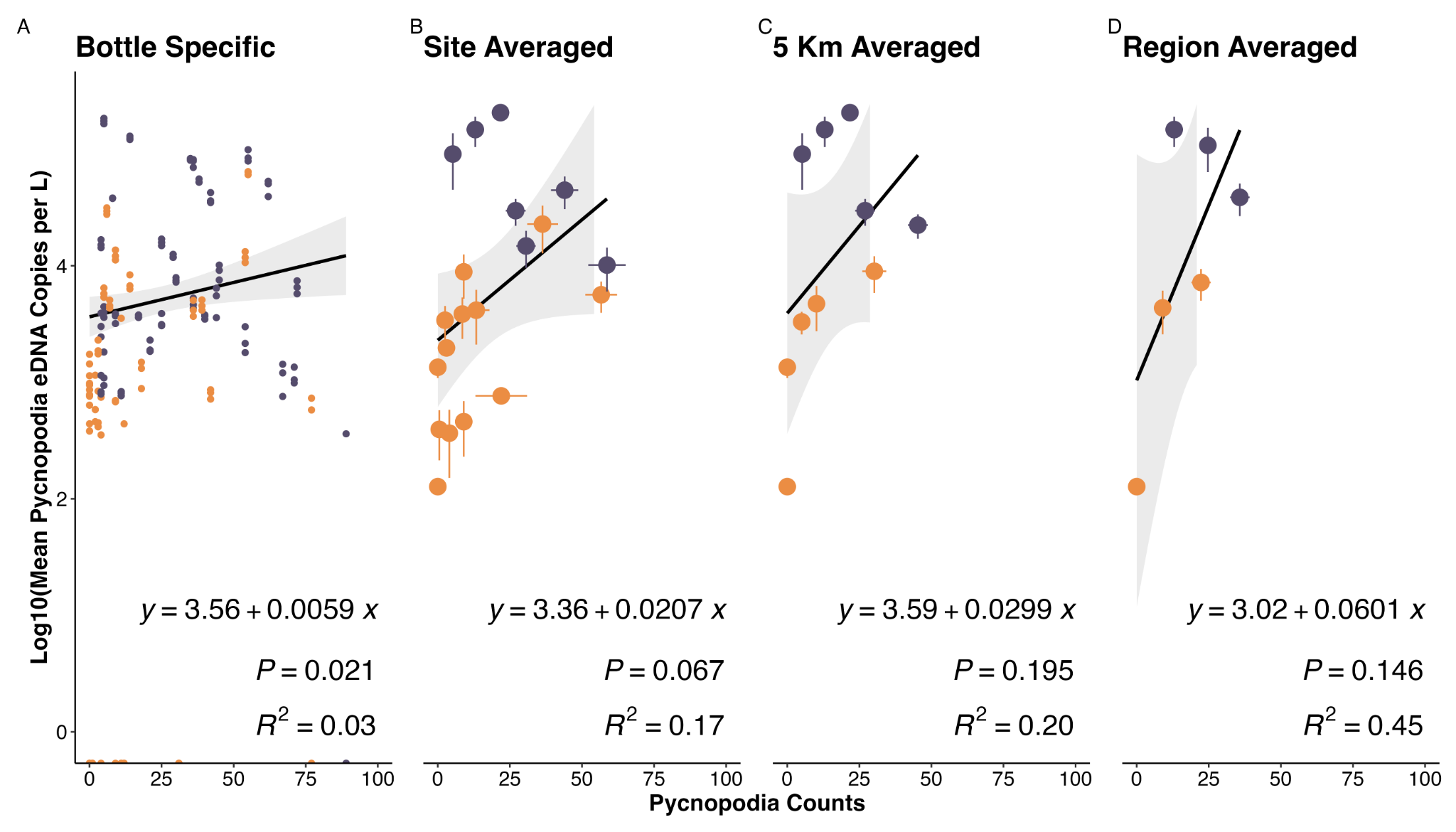

**Figure S14**. **Comparison of eDNA concentration with LoD filtering and underwater visual census based pycnopodia counts across increasing geographic scale.** Samples below LoD were set to zero. eDNA concentrations measured by ddPCR assay after log10 transformation. We find better alignment in eDNA and diver abundance estimates when averaging at larger scales as compared to point transect and bottle comparisons. We also find better alignment with eDNA biomass density estimates, which account for individual size (Figure 1) than with counts.
